# A contiguous *de novo* genome assembly of sugar beet EL10 (*Beta vulgaris* L.)

**DOI:** 10.1101/2020.09.15.298315

**Authors:** J. Mitchell (Mitch) McGrath, Andrew Funk, Paul Galewski, Shujun Ou, Belinda Townsend, Karen Davenport, Hajnalka Daligault, Shannon Johnson, Joyce Lee, Alex Hastie, Aude Darracq, Glenda Willems, Steve Barnes, Ivan Liachko, Shawn Sullivan, Sergey Koren, Adam Phillippy, Jie Wang, Tiffany Liu, Jane Pulman, Kevin Childs, Anastasia Yocum, Damian Fermin, Effie Mutasa-Göttgens, Piergiorgio Stevanato, Kazunori Taguchi, Kevin Dorn

**Author notes:** The author responsible for distribution of materials integral to the findings presented in this article in accordance with the policy described in the Instructions for Authors (www.plantcell.org) is: Mitch McGrath.

## Abstract

A contiguous assembly of the inbred ‘EL10’ sugar beet (*Beta vulgaris* ssp. *vulgaris*) genome was constructed using PacBio long read sequencing, BioNano optical mapping, Hi-C scaffolding, and Illumina short read error correction. The EL10.1 assembly was 540 Mb, of which 96.7% was contained in nine chromosome-sized pseudomolecules with lengths from 52 to 65 Mb, and 31 contigs with a median size of 282 kb that remained unassembled. Gene annotation incorporating RNAseq data and curated sequences via the MAKER annotation pipeline generated 24,255 gene models. Results indicated that the EL10.1 genome assembly is a contiguous genome assembly highly congruent with the published sugar beet reference genome. Gross duplicate gene analyses of EL10.1 revealed little large-scale intra-genome duplication. Reduced gene copy number for well-annotated gene families relative to other core eudicots was observed, especially for transcription factors. Variation in genome size in *B. vulgaris* was investigated by flow cytometry among 50 individuals drawn from EL10 progeny and three unrelated germplasm accessions, producing estimates from 633 to 875 Mb/1C. Read depth mapping with short-read whole genome sequences from other sugar beet germplasm suggested that relatively few regions of the sugar beet genome appeared associated with high-copy number variation.

## Introduction

Humans have used beet (*Beta vulgaris* spp. *vulgaris* L.) as early as the late Mesolithic, initially as leafy pot herb and for medicinal uses (Biancardi et al. 2012). It was not until the Middle Ages that the enlarged taproot was widely used as a vegetable. The origin of the enlarged taproot is not clear, but by the 18^th^ century beets were widely used as fodder and fueled the prelude to the Industrial Revolution in Europe. Sugar beet was selected from lower sucrose fodder beets (6-8% sucrose fresh weight) from the late 1700’s, with the first true sugar beet commercial varieties available by 1860 (Galon and Zallen 1998). Since then, improvements in sucrose content and processing quality have been continuous, resulting in an industry average in the US and Europe approaching 19% sucrose fresh weight (~75% dry weight). Breeding methods for sugar beet are applicable to the *B. vulgaris* vegetable crop types (table beet/ beet root and leafy chard) and fodder/ biofuel/ industrial chemical feedstock crop types (McGrath and Panella 2019, McGrath and Townsend 2015). Public sector sugar beet breeding today focuses generally on crop protection traits (Panella et al. 2008, 2015). The EL10.1 genome summarized here was recently interrogated for resistance gene signatures (Funk et al. 2018) and crop-type attributes (Galewski and McGrath 2020). An alternate assembly, EL10.2, is available but not as well characterized as EL10.1.

*Beta vulgaris* is a basal eudicot in the family Amaranthaceae (Caryophyllales) (Yang et al. 2015). Wild forms are found around European and Mediterranean coastlines and collectively classified as subspecies *maritima* (Biancardi et al. 2012, Andrello et al. 2016, 2017). There are no known barriers to cross-fertilization among beet crop and wild types, and the genomes of crop wild relatives are beginning to be described in detail (del Rio et al. 2019). Most *Beta vulgaris* types, and all characterized *maritima* types, are diploid. Chromosomes are morphologically similar at mitotic metaphase, and highly repetitive DNA sequences comprise ~60% or more of the beet genome (Flavell et al. 1974, Dohm 2014). Each chromosome shows different patterns of repeat-sequence distribution (Schmidt and Heslop-Harrison 1998, Paesold et al. 2012) supporting the notion that sugar beet genomes are true diploids (Halldén et al. 1998, Dohm et al. 2014). An ancient genome triplication appears to be shared with the basal asterid and rosid eudicot clades (Dohm et al. 2012). A uniform linkage group nomenclature was derived from Schondelmaier and Jung’s (1997) linkage group assignments and made more portable with SSR markers (McGrath et al. 2007). Extensive marker technologies remain proprietary within the commercial sugar beet breeding sector who supply hybrid seed to growers worldwide.

We seek to fill knowledge gaps in understanding of sugar beet traits by completing a genome framework for beet and then building crop genetic traits into the framework, focusing on crop quality and preservation traits. Creating highly contiguous genome assemblies is challenging, especially in plants due to the generally high-repetitive nature of portions of their genomes. Genome annotation is perhaps more challenging as expressed gene functions are generally predicted from relatively few physiologically-verified protein functions derived from unrelated plant taxa on the basis of nucleotide and amino acid sequence similarity. Improved approaches are becoming available and more commonly used (Jung et al. 2019). Many of these approaches were used in create the EL10 genome assemblies described here, including long-read length technologies which can span many (but not all) longer low-complexity repeat regions, optical mapping which can create larger scaffolds from long-read contig assemblies, and Hi-C which can link together scaffolds across the genome into chromosome-sized scaffolds. Highly contiguous assemblies exhibit the full organization of hereditary material, thus little uncertainty of position and distribution of genetic markers, for instance, allows closer focus on any region of the genome.

Scaffolds of the EL10 assemblies show high concordance with genetic maps and the RefBeet genome sequence (Dohm et al. 2014), which is an excellent but fragmented genome sequence assembled using first- and second-generation sequencing technologies. The current work from less fragmented assemblies used here provide a more comprehensive picture of genome size variability of sugar beet which surprisingly varies extensively each generation, and global changes in repeat sequence depth and coverage within and between sugar beet inbreds and breeding populations. Genome fluidity generates mutations, and assessing whether these recombinational mutation events are useful for sugar beet improvement, or simply a hinderance, can be investigated. Further, gene number in beet appears to be uniformly diminished relative to other eudicots, at least for gene classifiers that are shared among representative angiosperm genomes. Contiguous genome assemblies will allow routine inter-cultivar comparisons between accessions that vary for important traits, and thus help deduce casual from associated genomic features influencing a trait of interest, general performance, or otherwise suggest that an association is merely an historical coincidence of shared parentage and breeding.

## Results

A five-generation inbred genome of the sugar beet ‘C869’ (PI 628755) was released as a genetic stock ‘EL10’ (PI 689015). C869 is the common seed parent for East Lansing recombinant inbred populations (McGrath et al. 2005). Five plants from one inbred family showing no gross phenotypic differences and no polymorphism among 24 selected unlinked SSR markers (McGrath et al. 2007) were chosen for nuclei isolation, long read sequencing, and assembly. The resulting assembly, using only one of the five plants, was scaffolded via opto-physical mapping, and the two assemblies described here share this common backbone. Hi-C scaffolding was largely able to reduce the number of scaffolds to the haploid chromosome number in beet (n=9), with assembly EL10.2 slightly improved in contiguity over assembly EL10.1. Insights described below pertain to EL10.1 since the annotation of EL10.2 is on-going.

### Sequencing and Assembly

High molecular weight DNA was isolated from intact gel-embedded nuclei of true leaves from young seedlings and pooled from the five inbred plants for long-read sequencing using standard protocols for BAC library construction (Amplicon Express, Pullman, WA). Eighty-six PacBio SMRT cells yielded 79.3-fold coverage (58,655 Mb) of the *circa* 750 Mb *Beta vulgaris* genome size (see below). The Falcon Assembler (version 0.2.2) was used to assemble long reads (Table 1), initialized with reads exceeding 40 kb in length. The Falcon assembly resulted in 938 primary contigs, 70.9% with a length greater than 100,000 nucleotides and a total length of 562.76 Mb (Table 2). Both assembly versions (EL10.1 and EL10.2) relied on this intermediate assembly (e.g. SBJ_80X_BN, Table 2). G+C content was similar between EL10 and RefBeet contigs (35.8% vs. 36.1%, respectively).

**Table 1:**
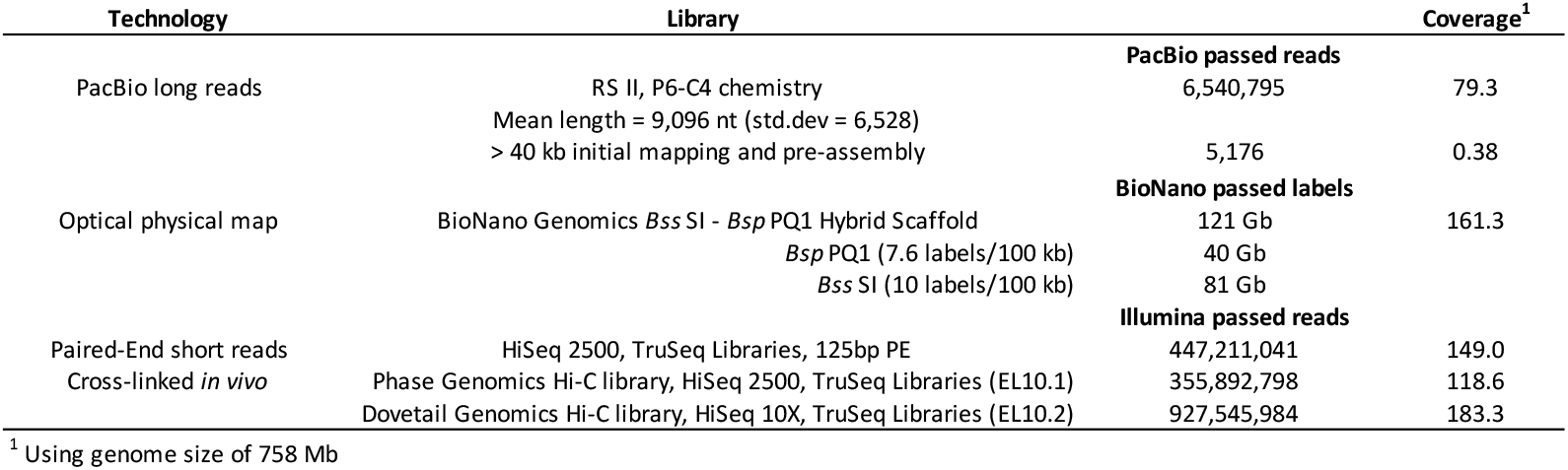
Sequence inputs and metrics used in construction of EL10.1.

**Table 2:** Assembly metrics for EL10.1 and sequence assembly iterations.

| Assembly by input and method | Name | # Contigs | % Scaffolded | Total size | N50 | NG50<sup>1</sup> | % >100 kb | # Scaffolds | Total size | N50 | NG50<sup>1</sup> | %N | Coverage %<sup>1</sup> |
| --- | --- | --- | --- | --- | --- | --- | --- | --- | --- | --- | --- | --- | --- |
|  |  |  |  | (x 1,000 nt) |  |  |  |  | (x 1,000 nt) |  |  |  |  |
| RefBeet 1.2 (Dohm et al. 2014) | RefBeet | 60,051 | 93.7 | 517,882 | 43.8 | nd | 1.0 | 40,508 | 566,571 | 2,013 | nd | 8.60 | nd |
| EL10.1 PacBio | SBJ_80X | 938 | na | 562,760 | 1,394 | 1,228 | 70.9 | 938 | 562,760 | 1,394 | 1,228 | 0.00 | 89.6 |
| EL10.1 PacBio BioNano | SBJ_80X_BN | 2,983 | 99.2 | 533,042 | 1,340 | 1,093 | 21.5 | 86 | 566,848 | 12,513 | 10,655 | 5.90 | 90.3 |
| EL10.1 PacBio BioNano Hi-C | EL10.1 | 364 | 96.2 | 540,479 | 2,701 | 2,335 | 96.7 | 40 | 540,537 | 57,939 | 57,353 | 0.01 | 86.1 |
|  | Chromosome_1 | 47 | 100 | 58,076 | 2,421 | nd | 100.0 | 1 | 58,086 | na | nd | 0.02 | 9.2 |
|  | Chromosome_2 | 30 | 100 | 54,968 | 2,834 | nd | 96.7 | 1 | 54,972 | na | nd | 0.01 | 9.2 |
|  | Chromosome_3 | 22 | 100 | 54,096 | 3,728 | nd | 100.0 | 1 | 54,100 | na | nd | 0.01 | 8.6 |
|  | Chromosome_4 | 47 | 100 | 61,154 | 2,396 | nd | 97.9 | 1 | 61,163 | na | nd | 0.01 | 9.7 |
|  | Chromosome_5 | 30 | 100 | 59,218 | 3,579 | nd | 93.3 | 1 | 59,225 | na | nd | 0.01 | 9.4 |
|  | Chromosome_6 | 52 | 100 | 65,091 | 2,381 | nd | 98.1 | 1 | 65,097 | na | nd | 0.01 | 10.4 |
|  | Chromosome_7 | 40 | 100 | 57,345 | 2,831 | nd | 95.0 | 1 | 57,354 | na | nd | 0.02 | 9.1 |
|  | Chromosome_8 | 37 | 100 | 57,932 | 2,335 | nd | 97.3 | 1 | 57,939 | na | nd | 0.01 | 9.2 |
|  | Chromosome_9 | 28 | 100 | 52,176 | 2,382 | nd | 100.0 | 1 | 52,180 | na | nd | 0.01 | 8.3 |
|  | Unplaced Scaffolds | 31 | 0 | 20,421 | 1,679 | nd | 87.1 | 31 | 20,421 | na | nd | 0.00 | 3.3 |
| EL10.2 PacBio BioNano Hi-C | EL10.2 | 3,098 | 99.9 | 533,041 | 1,283 | nd | 21.1 | 18 | 567,031 | 61,792 | nd | 5.99 | 90.3 |
|  | Chromosome_1 | 505 | 100 | 58,689 | 1,093 | nd | 16.0 | 1 | 64,154 | na | nd | 8.52 | 10.2 |
|  | Chromosome_2 | 253 | 100 | 53,946 | 1,566 | nd | 21.3 | 1 | 56,769 | na | nd | 4.97 | 9.0 |
|  | Chromosome_3 | 181 | 100 | 54,788 | 1,907 | nd | 28.7 | 1 | 57,123 | na | nd | 4.00 | 9.1 |
|  | Chromosome_4 | 406 | 100 | 61,919 | 1,225 | nd | 20.0 | 1 | 66,143 | na | nd | 6.39 | 10.5 |
|  | Chromosome_5 | 276 | 100 | 64,991 | 1,635 | nd | 21.4 | 1 | 67,720 | na | nd | 4.03 | 10.8 |
|  | Chromosome_6 | 423 | 100 | 68,152 | 1,004 | nd | 23.6 | 1 | 72,250 | na | nd | 5.67 | 11.5 |
|  | Chromosome_7 | 349 | 100 | 57,001 | 1,144 | nd | 22.6 | 1 | 60,906 | na | nd | 6.41 | 9.7 |
|  | Chromosome_8 | 245 | 100 | 59,411 | 1,155 | nd | 31.8 | 1 | 61,792 | na | nd | 3.85 | 9.8 |
|  | Chromosome_9 | 308 | 100 | 51,533 | 1,527 | nd | 18.8 | 1 | 55,602 | na | nd | 7.32 | 8.9 |
|  | Unplaced Scaffolds | 152 | 97.4 | 2,613 | 252 | nd | 0.0 | 9 | 4,573 | na | nd | 42.86 | 0.7 |

<sup>1</sup> Based on 628 Mb Physical Map

Scaffolding the Falcon assembly with a BioNano two-enzyme (*BspQI* and *Bss*SI) sequential hybrid optical (physical) map resulted in substantial improvement. The *Bsp*QI optical map was generated from 141,462 molecules with an average length of 285 kb and labelled to an average density of 11.8 sites kb^-1^, and the *Bss*SI optical map was generated from 270,071 molecules, also with an average length of 285 kb, labelled to a density of 7.7 sites kb^-1^. Optical maps were aligned to PacBio Falcon contigs and the resulting *Bsp*QI and *Bss*SI map lengths were 628 Mb and 590 MB with N50 contig sizes of 1.99 Mb and 1.21 Mb, respectively. After merging PacBio, *BspQI*, and *Bss*SI contigs, the final hybrid genome map consisted of 86 scaffolds with a total length of 566.8 Mb, and an N50 of 12.5 MB (Table 2).

Hi-C Proximity Guided Assembly, using selfed progeny of the individual that was optically mapped (see below), was applied to the merged PacBio/BioNano assembly (e.g. SBJ_80X_BN). This assembly was polished and gap-filled using a combination of approaches (PBJelly, Arrow, and Pilon; following Bickhart et al. 2017). The resulting 540.5 Mb assembly consisted of 9 chromosome-sized scaffolds, numbered via Butterfass chromosome nomenclature (Butterfass 1964), and 31 unscaffolded contigs. These comprise the genome assembly version described here, EL10.1. The 9 chromosome-sized scaffolds (designated Chromosomes below) were relatively similar in size (mean = 57.8 Mb, std. dev. = 3.9 Mb) (Table 2). Chromosomally-unscaffolded contigs (n=31, hereafter referred to as Scaffolds) represented 3.9% of the final EL10.1. A second assembly (EL10.2) was created using a second set of Hi-C reads and the default Dovetail Genomics HiRise assembly pipeline, starting from the PacBio-BioNano assembly SBJ_80X_BN (Table 2).

The EL10.1 genome assembly was not corrected for the differences described below, however EL10.1 has been used in at least two publications (Funk et al. 2018, Galewski and McGrath 2020), and therefore is important to document. Differences between assemblies have not been fully annotated and further improvement of the EL10 assembly is likely, thus, the EL10.2 assembly is being reported here simply as a publicly available resource for community inspection, assessment, and basis for refinement. Initial assessment suggested the EL10.2 assembly appeared to have resolved the major assembly-associated inversions on Chromosomes 7 and 9 (see below), as well as placed the 31 Scaffolds into the larger whole genome chromosome context (Figure 2), where many unplaced Scaffolds of EL10.1 appeared to be better placed within the context of Chromosome 5 in EL10.2.

**Figure 1:**
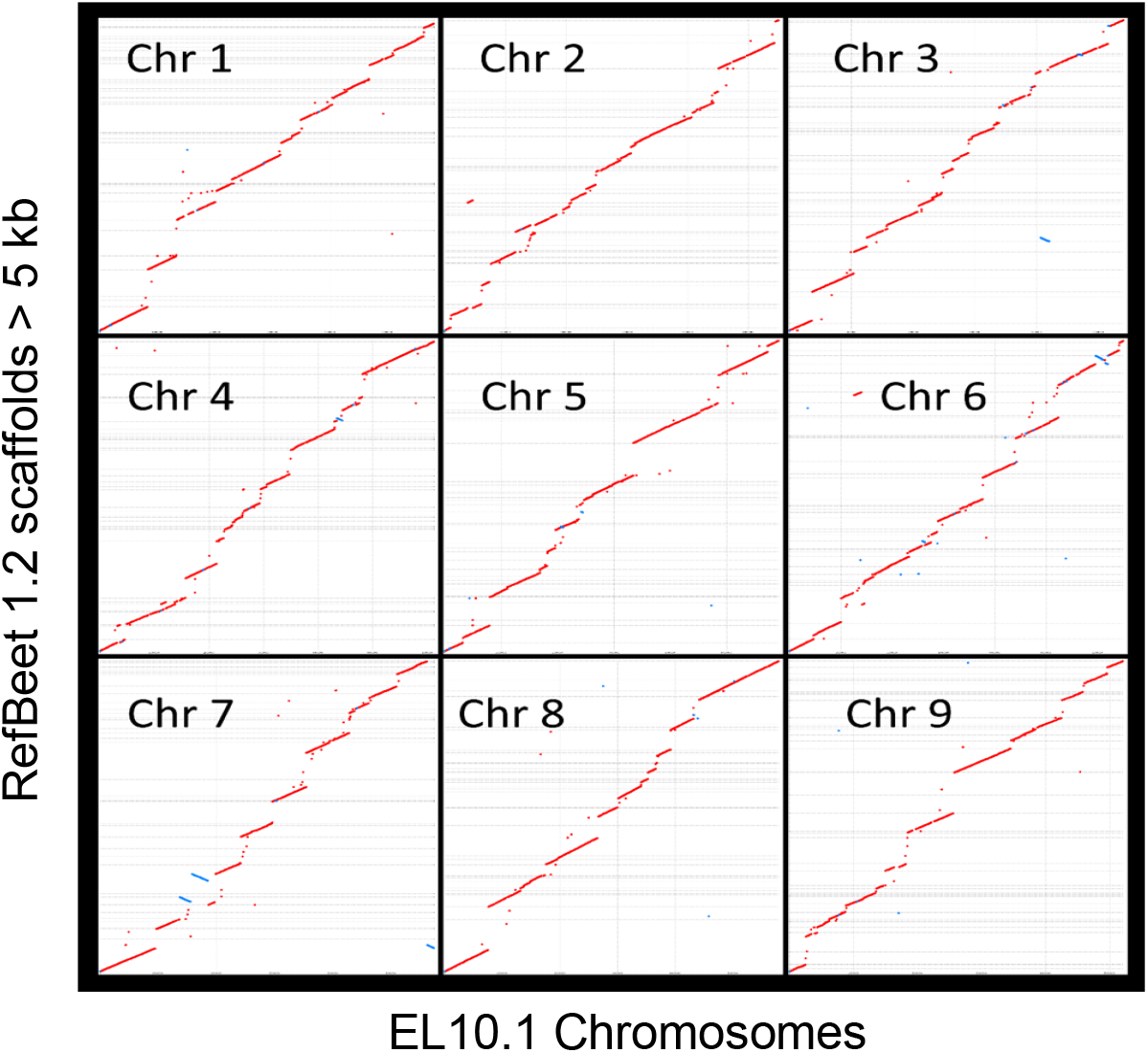
Chromosome alignment of the EL10 assembly (x-axis) versus RefBeet-1.2 assembly (y-axis) by EL10.1 Chromosome. Alignments less than 5 kb in length were removed before plotting. Alignments with matching orientation are shown in red, inversions are show in blue. Unassembled RefBeet regions are indicated by gaps.

**Figure 2:**
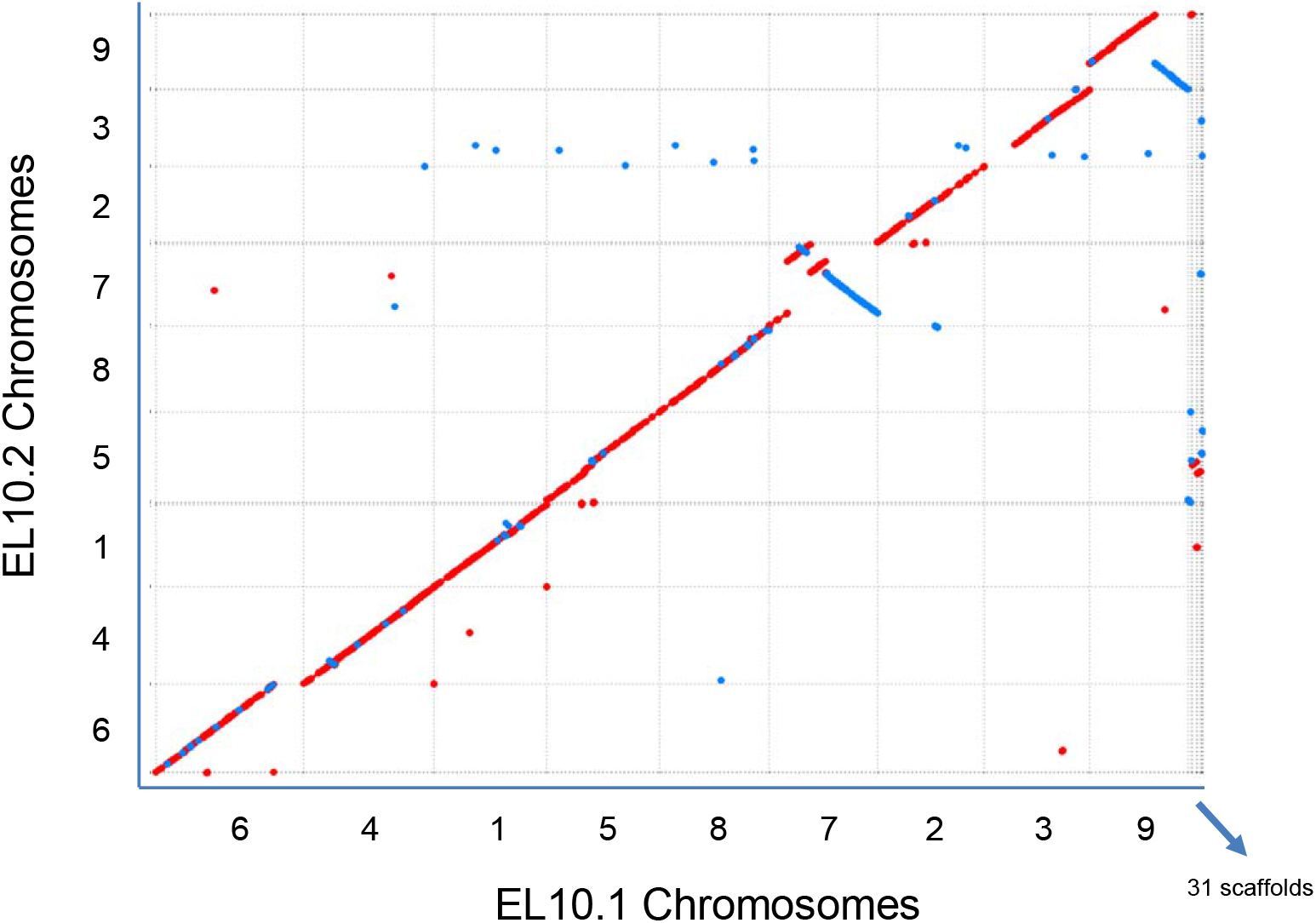
Comparison of contiguity between EL10.1 and EL10.2 genome assemblies. Alignments with matching orientation are shown in red, inversions are show in blue.

### Assessment

No complete chloroplast or mitochondrial genomes were incorporated into the EL10.1 assembly, although fragments of both plastid genomes were detected in the EL10.1 assembly. The position of RefBeet 1.2 scaffolds were determined for EL10.1 Chromosomes (Figure 1). Contigs > 5 kb in length were largely colinear between the two assemblies. Two small inverted-orientation contigs were evident on Chromosome 7, as were small inverted (e.g. Chromosome 6) and misplaced segments (e.g. Chromosomes 3 and 7). RefBeet 1.2 was anchored with genetic markers (Dohm et al. 2014), and 345 of these with 100% match identity across 75 nt or greater were placed in concordant order on the EL10.1 assembly. In addition, 3,279 proprietary SNP markers from the SESVanderhave (Tienen, Belgium) molecular marker genetic map were placed to the EL10.1 assembly. Most marker orders were highly concordant. However, a third of the mapped markers were inverted on Chromosome 9, and a complex rearrangement involving 40% of markers was evident on Chromosome 7 (mapped inversions; Table 3). Genetic markers also added 9 Scaffolds to five Chromosomes (mapped integrations; Table 4). Genetic markers used to orient the cytogenetic map (Paesold et al. 2012) also aligned with the EL10.1 assembly. Chromosomes 1 and 3 were cytogenetically congruent with their North-South orientation, and the rest were reversed relative to the orientations given in that publication. Scaffold 5 was located to the South end of cytogenetic Chromosome 5 (Table 5), consistent with SESVanderhave marker data (Table 4).

**Table 3:** Inversions in the EL10.1 genome assembly assessed using genetic markers.

| EL10.1 | Position (Mb) | SES Marker<br>Position |
| --- | --- | --- |
| Chromosome_7 | 0.2 | 7 |
| Chromosome_7 | 9.5 | 45 |
| Chromosome_7 | 10.3 | 30 |
| Chromosome_7 | 14.1 | 28 |
| Chromosome_7 | 15.8 | 18 |
| Chromosome_7 | 19.7 | 27 |
| Chromosome_7 | 19.9 | 18 |
| Chromosome_7 | 21.8 | 3 |
| Chromosome_7 | 24.8 | 32 |
| Chromosome_7 | 57.3 | 44 |
| Chromosome_9 | 3.1 | 67 |
| Chromosome_9 | 25.2 | 50 |
| Chromosome_9 | 25.8 | 8 |
| Chromosome_9 | 34.5 | 50 |
| Chromosome_9 | 41.3 | 71 |
| Chromosome_9 | 50.6 | 74 |

**Table 4:** Co-locations of Scaffolds and Chromosomes deduced by genetically mapped markers.

| EL10.1<br>Chromosome | EL10.1<br>Scaffold | Orientation | Bin Position (genomic<br>coordinates) |
| --- | --- | --- | --- |
| Chromosome_1 | Scaffold_07 | reverse | 58,086,001 - end |
| Chromosome_2 | Scaffold_19 | unknown | 30,001,550 - 30,051,550 |
| Chromosome_3 | Scaffold_03 | reverse | 27,470,050 - 27,610,050 |
| Chromosome_5 | Scaffold_05 | reverse | end - 216,554 |
| Chromosome_5 | Scaffold_04 | forward | 19,422,050 - 19,462,050 |
| Chromosome_5 | Scaffold_01 | forward | 24,502,050 - 24,632,050 |
| Chromosome_5 | Scaffold_14 | reverse | 45,192,050 - 45,212,050 |
| Chromosome_6 | Scaffold_02 | forward | 14,450,050 - 14,610,050 |
| Chromosome_6 | Scaffold_08 | forward | 61,492,050 - 61,552,050 |

**Table 5:**
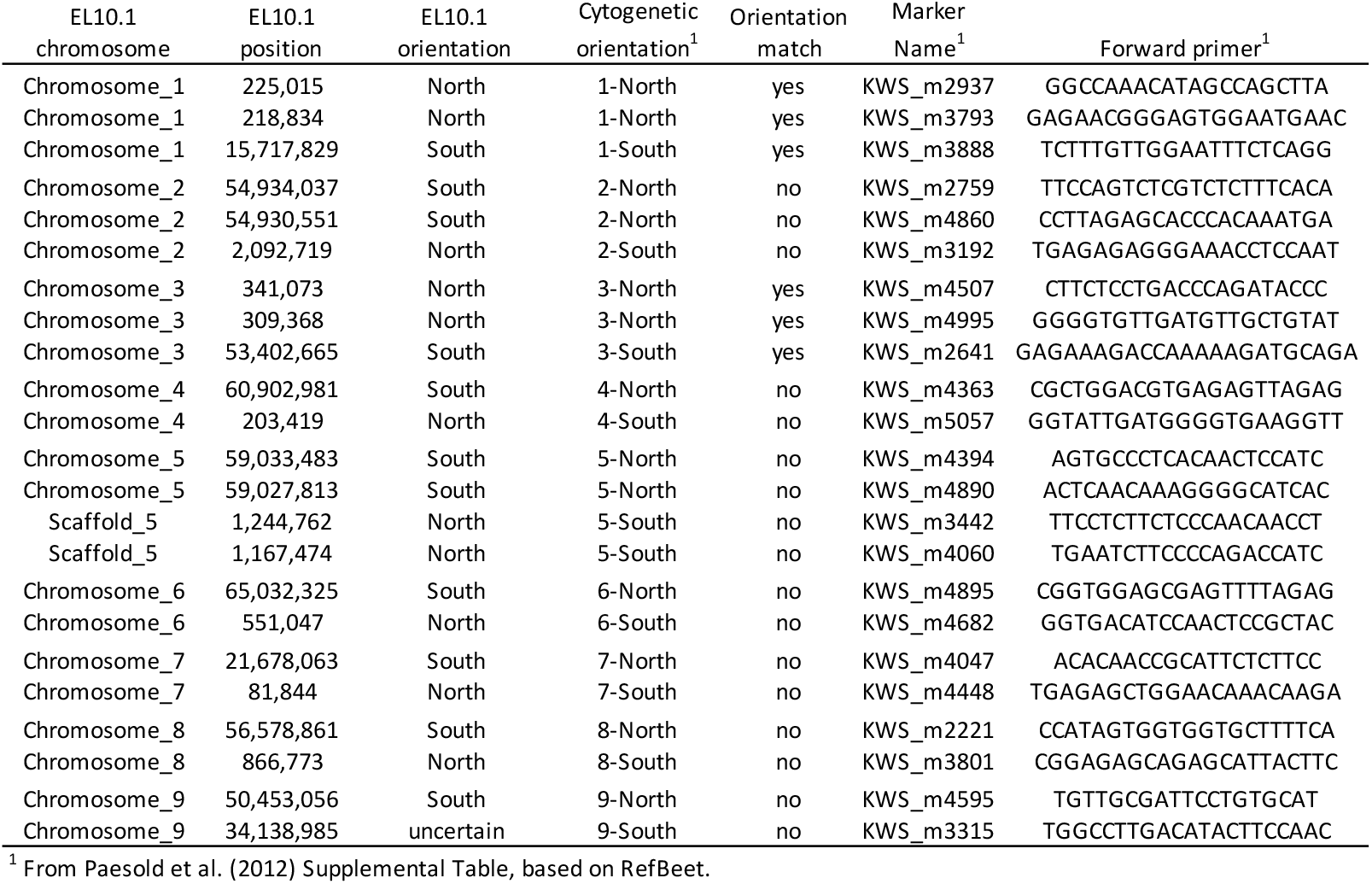
Orientation of EL10.1 Chromosomes relative to the cytogenetic map of Paesold et al. (2012).

### Annotation

Of note, the EL10.1 assembly contained the entire first linkage group described in beet (Keller 1936), the *R-Y-B* linkage group on Chromosome 2. Each of these genes has been recently cloned (*R*, for the red alkaloid betalain synthesis by a novel cytochrome P450; Hatlestad et al. 2012), *Y*, a Myb transcription factor required for production of red color (Hatlestad et al. 2014), and *B* for the bolting gene which determines annual or biennial life habit; Pin et al. 2010, 2012). Both the direction and the distance agree with published genetic map intervals, and the EL10.1 assembly indicates that the bolting gene is physically located proximal towards the centromere and the color genes are more distal (Table 6).

**Table 6:** The Y-R-B linkage group in the EL10.1 genome assembly.

| Gene | ID | MAKER Inferred Annotation | EL10.1_start..stop (strand) | cM<sup>1</sup> | Mb | Mb/cM |
| --- | --- | --- | --- | --- | --- | --- |
| *Y* | EL10Ac2g04466 | MYB114 | 49,675,759..49,679,064 (plus) | 7.4 | 2.4 | 0.32 |
| *R* | EL10Ac2g04268 | Geraniol 8-hydroxylase | 47,304,905..47,309,543 (minus) | 17.3 | 14.7 | 0.85 |
| *B* | EL10Ac2g03535 | Response regulator-like PRR73 | 32,610,511..32,619,244 (minus) |  |  |  |

<sup>1</sup> From Abe (1993), Goldman and Austin (2000)

Results from the MAKER annotation pipeline (Holt and Yandell 2011) conservatively predicted 24,255 proteins, numerically 88.5% of the 27,421 predicted in RefBeet (Dohm et al. 2014). For functional annotation, three sources were used, in the priority: 1) UniProt, 2) Pfam-A, and 3) Uniref90. If no functional annotation was found in these three highly curated sets, predicted proteins were assigned to the class of ‘hypothetical’ proteins. Gene model completeness was checked using BUSCO (Table 7) (Simão et al. 2015). A higher proportion of missing BUSCO’s was seen in EL10.1 than either RefBeet 1.1 or Arabidopsis. Overall, protein coding gene predictions covered a relatively small proportion of the assembled EL10.1 genome (39,161,207 nt; 7.2%). GC content of predicted coding genes was marginally higher than that of the whole genome (41.1% vs. 35.8%, respectively). Predicted proteins were named using the underlined characters in the key: <u>EL10</u> / annotation version <u>A</u> / <u>c</u>hromosome or <u>s</u>caffold number / genomic in origin / a sequential number / and appended with <u>.1</u> to signify that only one isoform was considered at this level of analysis (e.g. EL10Ac7g16740.1).

**Table 7:** Gene models detected via Benchmarking Universal Single-Copy Orthologs (BUSCO).

| BUSCO | EL10.1 | RefBeet 1.1 | TAIR 10 |
| --- | --- | --- | --- |
| Complete | 1,251 | 1,302 | 1,414 |
| Complete and single-copy | 1,223 | 1,268 | 1,401 |
| Complete and duplicated | 28 | 34 | 13 |
| Fragmented | 36 | 37 | 7 |
| Missing | 153 | 101 | 19 |
| Total groups searched | 1,440 | 1,440 | 1,440 |
| % Missing | 10.6 | 7.0 | 1.3 |

The number of MAKER annotations ascribed across Chromosomes of EL10.1 was relatively consistent (mean = 2,559, stdev = 173.8), but highly variable between Scaffolds (mean = 44.0, stdev = 47.6) (excluding Scaffolds 23, 29, 30, and 31 for which no gene models were predicted) (Table 8). A total of 3,940 gene models had no functional annotation among curated comparative databases (and thus were designated hypothetical), and these were also evenly distributed among Chromosomes but not necessarily Scaffolds (Table 8). Fewer than 55% of gene models were considered unique in the sense their curated-database annotations only occurred once in the list of gene models (Table 8), and thus, at this level of analysis, more than 45% of predicted genes could be members of gene families.

**Table 8:** Distribution of MAKER annotations across the EL10.1 genome assembly.

| Location | # genes | # hypothetical | % | Unique | % |
| --- | --- | --- | --- | --- | --- |
| Total | 24,255 | 3,940 | 16.24 | 13,220 | 54.50 |
| Chromosome 1 | 2,393 | 354 | 14.79 | 1,375 | 57.46 |
| Chromosome 2 | 2,525 | 383 | 15.17 | 1,379 | 54.61 |
| Chromosome 3 | 2,574 | 372 | 14.45 | 1,458 | 56.64 |
| Chromosome 4 | 2,903 | 463 | 15.95 | 1,410 | 48.57 |
| Chromosome 5 | 2,687 | 422 | 15.71 | 1,452 | 54.04 |
| Chromosome 6 | 2,681 | 446 | 16.64 | 1,478 | 55.13 |
| Chromosome 7 | 2,487 | 385 | 15.48 | 1,380 | 55.49 |
| Chromosome 8 | 2,391 | 353 | 14.76 | 1,316 | 55.04 |
| Chromosome 9 | 2,387 | 338 | 14.16 | 1,320 | 55.30 |
| Scaffold 1 | 117 | 54 | 46.15 | 55 | 47.01 |
| Scaffold 2 | 141 | 60 | 42.55 | 65 | 46.10 |
| Scaffold 3 | 87 | 39 | 44.83 | 48 | 55.17 |
| Scaffold 4 | 94 | 38 | 40.43 | 46 | 48.94 |
| Scaffold 5 | 162 | 22 | 13.58 | 121 | 74.69 |
| Scaffold 6 | 79 | 28 | 35.44 | 36 | 45.57 |
| Scaffold 7 | 119 | 50 | 42.02 | 54 | 45.38 |
| Scaffold 8 | 91 | 15 | 16.48 | 66 | 72.53 |
| Scaffold 9 | 35 | 15 | 42.86 | 16 | 45.71 |
| Scaffold 10 | 25 | 10 | 40.00 | 15 | 60.00 |
| Scaffold 11 | 19 | 12 | 63.16 | 3 | 15.79 |
| Scaffold 12 | 46 | 10 | 21.74 | 33 | 71.74 |
| Scaffold 13 | 21 | 10 | 47.62 | 11 | 52.38 |
| Scaffold 14 | 30 | 6 | 20.00 | 15 | 50.00 |
| Scaffold 15 | 52 | 10 | 19.23 | 20 | 38.46 |
| Scaffold 16 | 5 | 3 | 60.00 | 2 | 40.00 |
| Scaffold 17 | 3 | 1 | 33.33 | 1 | 33.33 |
| Scaffold 18 | 15 | 2 | 13.33 | 9 | 60.00 |
| Scaffold 19 | 30 | 17 | 56.67 | 5 | 16.67 |
| Scaffold 20 | 16 | 9 | 56.25 | 7 | 43.75 |
| Scaffold 21 | 5 | 3 | 60.00 | 0 | 0.00 |
| Scaffold 22 | 10 | 4 | 40.00 | 6 | 60.00 |
| Scaffold 24 | 2 | 1 | 50.00 | 2 | 100.00 |
| Scaffold 25 | 1 | 0 | 0.00 | 1 | 100.00 |
| Scaffold 26 | 12 | 3 | 25.00 | 9 | 75.00 |
| Scaffold 27 | 4 | 2 | 50.00 | 0 | 0.00 |
| Scaffold 28 | 6 | 0 | 0.00 | 6 | 100.00 |
| Chromosome mean | 2,558.7 | 390.7 |  | 1,396.4 |  |
| Chromosome stdev | 173.8 | 43.7 |  | 58.2 |  |
| Scaffold mean | 43.8 | 15.1 |  | 23.3 |  |
| Scaffold stdev | 47.6 | 17.5 |  | 28.8 |  |

Self-synteny of MAKER gene models with the EL10.1 genome sequence was explored using the CoGe SynMap platform (Lyons et al. 2008). Few internal syntenies were detected. Mean copy number of the 2,327 discovered tandem gene models was 2.82 (stdev = 1.96), and 65.8% of these tandem duplications were two copies. For syntenic regions with at least 5 matches in a span of 20 gene models (encompassing 1,858 genes in 268 synteny blocks), average Kn/Ks values were all less than 1, suggesting stabilizing selection for genes in these blocks. For individual gene pairs, only five gene pairs had Kn/Ks values >1 (suggesting diversifying selection) but only two of these pairs had interpretable annotations. EL10Ac6g14284.1 & EL10Ac9g20883.1 were predicted as Clathrin heavy chain 2 genes (i.e., vesicle trafficking) and EL10Ac1g01568.1 & EL10Ac5g12109.1 were predicted SET Domain Protein genes (i.e., chromatin structure modulation).

A comparative gene annotation perspective was gained using the MapMan4 ontology of plant proteomes (Schwacke et al. 2019). EL10.1 MAKER gene models were placed in 99.6% of 4,145 ontologies assigned to one of 28 ‘bins’ (infrequently allowing for assignment to more than one bin), organized in a hierarchal, conceptual, plant-specific context (e.g., Photosynthesis, Cell cycle, Hormones, etc.). Where possible, each bin resolves to a gene from a high-quality genome assembly in the Mercator4 web implementation of MapMan4. Specific comparisons for each of the 4,127 EL10.1 occupied terminal, termed ‘leaf’, bins were made with five other angiosperms (e.g., *Arabidopsis thaliana, Oryza sativa, Brachypodium distachyon, Solanum lycopersicum*, and *Manihot esculenta*). Most EL10.1 predicted proteins in the found set were placed in one (or more) MapMan4 leaf bins (Table 9). Since the MapMan4 ontology is hierarchal, the number of genes in each leaf bin was averaged for all five angiosperms, and compared to EL10.1. Surprisingly, the number of genes in the EL10.1 gene set was 69% that of the average of five angiosperms (Table 9).

**Table 9:**
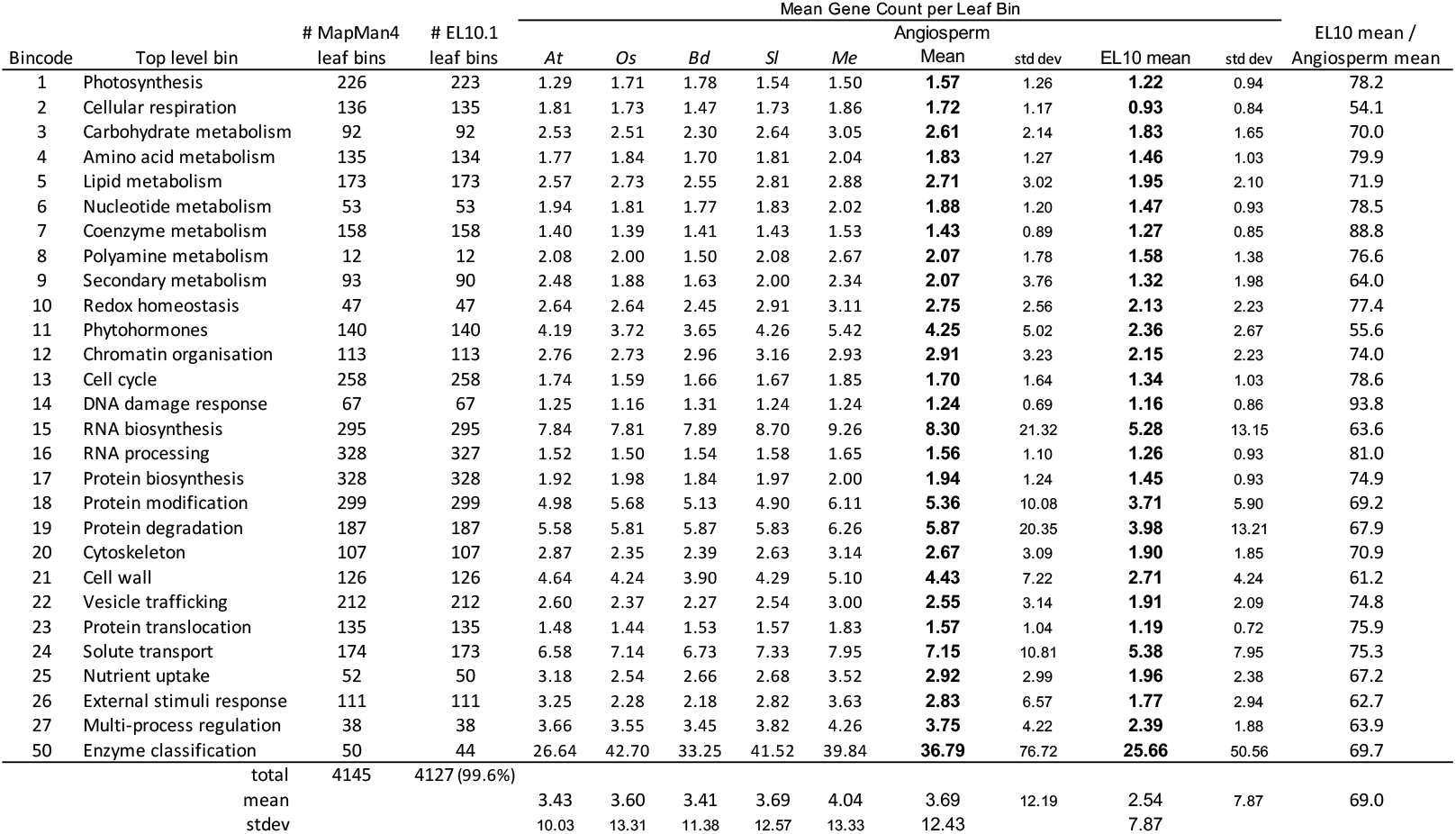
Comparison of MapMan4 first-order functional classifications for EL10.1 and four other Tracheophytes [*Oryza sativa (Os) Brachypodium distachyon (Bd), Arabidopsis thaliana (At), Solanum lycopersicum (Sl), and Manihot esculenta (Me)*].

Enrichment analysis can shed light on biological processes that may have assumed greater or lesser importance in the evolutionary success of a lineage. Given the general reduced gene copy number in EL10.1, genes whose copy number equaled or exceeded the mean of five angiosperms were tentatively considered as enriched, and those that were substantially lower than the overall mean of EL10.1 were considered as reduced. EL10.1 appeared particularly depauperate in at least two top-level ontologies; Cellular respiration (Bincode 2) and Phytohormones (Bincode 11) (Table 9). Equal or over-represented ontologies included DNA Damage Response (Bincode 14) and Coenzyme metabolism (Bincode 7) (Table 9).

Proteome content of the five averaged angiosperms relative to EL10.1 was gauged for missing members, which could suggest regions in EL10.1 that were not assembled, genes that were not annotated, or perhaps reflect biological divergence or biochemical alternatives that beet followed during its evolution. Not detected in EL10.1 were 154 genes that were present in at least one copy in each of the five angiosperms. Missing annotations were assignable across all 28 top-level bins, with the exception of Bincode 8 (Polyamine metabolism) (Table 10). In this set, mean copy number was low (1.6 genes per leaf bin) in the five taxa, and failure to assemble or annotate a low-copy number genes in EL10.1 was possible. However, in 12 cases, each of the five other plants had small gene families (mean copy number = 3.7 genes per family) but no EL10.1 homologue was annotated, which seemed less probable that all would have been missed during assembly and annotation, thus their functions in beet may have been dispensable, their genes diverged, or their functions assumed by other genes.

**Table 10:**
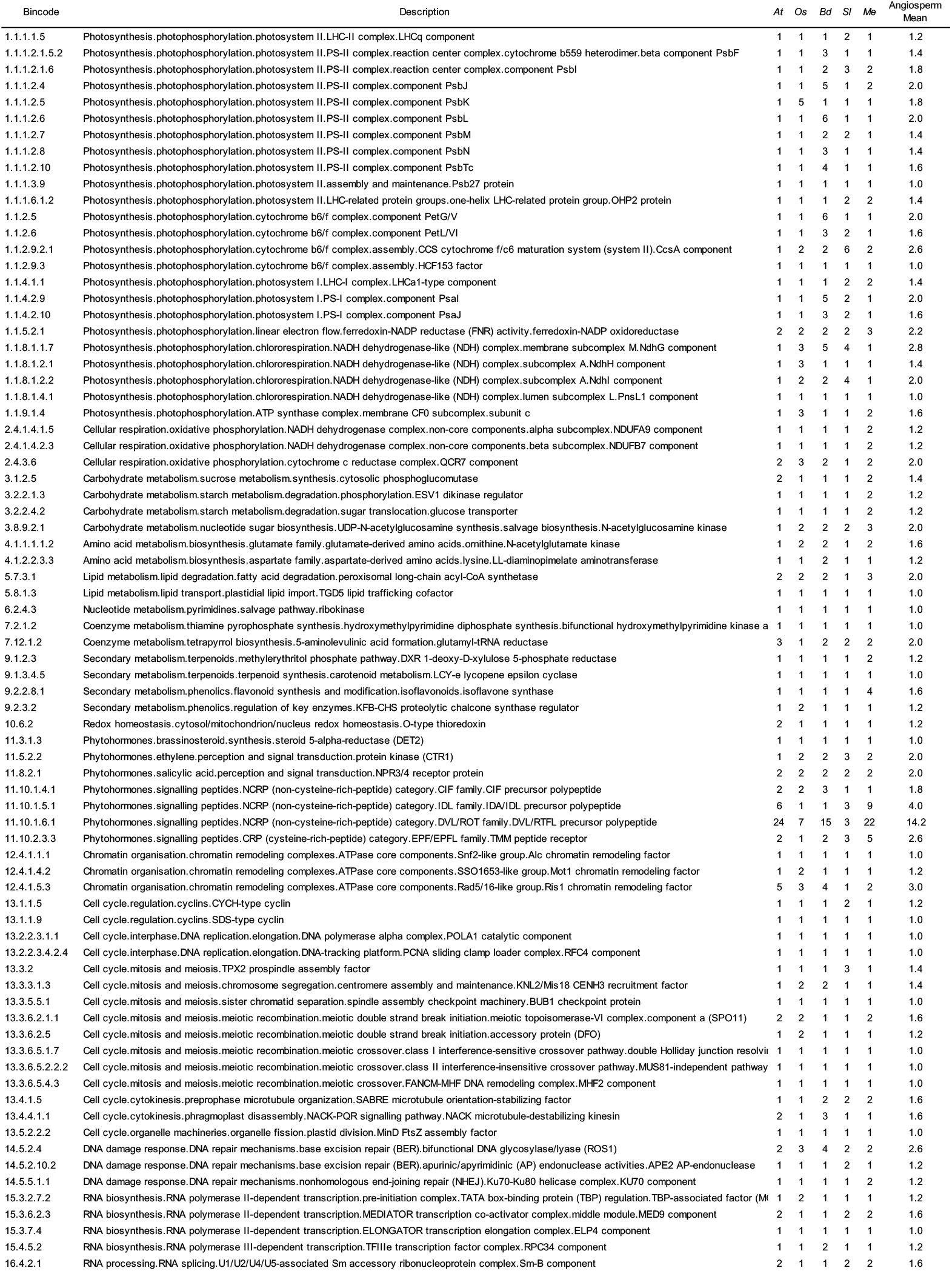
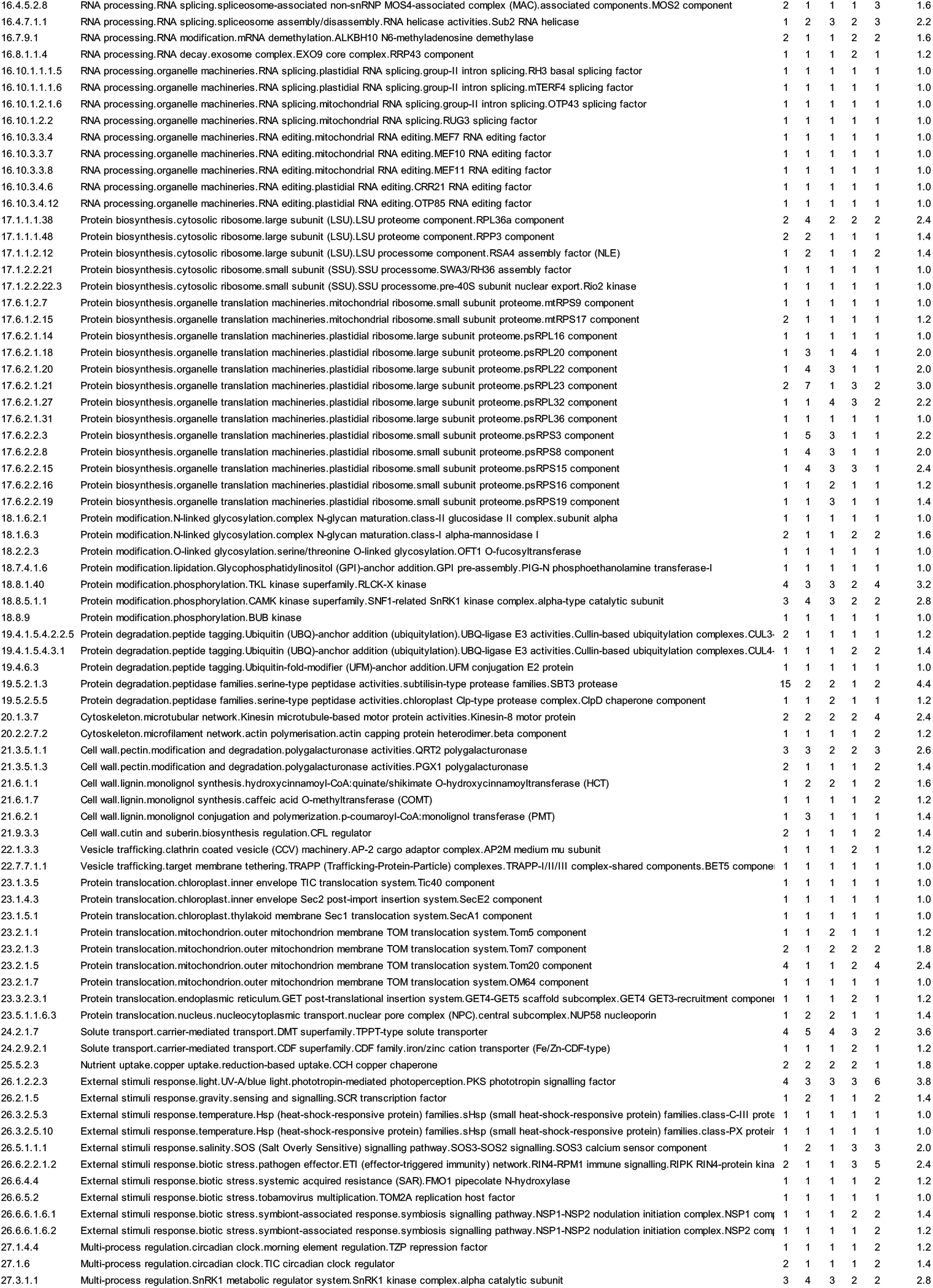
Comparison of MapMan4 leaf bins where no gene was predicted by MAKER in EL10.1 and five Tracheophytes [*Oryza sativa (Os) Brachypodium distachyon (Bd), Arabidopsis thaliana (At), Solanum lycopersicum (Sl), and Manihot esculenta (Me)*].

Remaining annotations were surveyed for potential biological interest, but not exhaustively evaluated (Table 11). Under-represented genes in ‘Cell wall’ (Bincode 21) included those involved with hemicellulose, lignin, cutin, and suberin metabolism, as might be expected from selection for a mechanically-sliced root crop for sucrose extraction (e.g., less knife wear during processing, which is a trait that has not necessarily been under conscious selection).

**Table 11:**
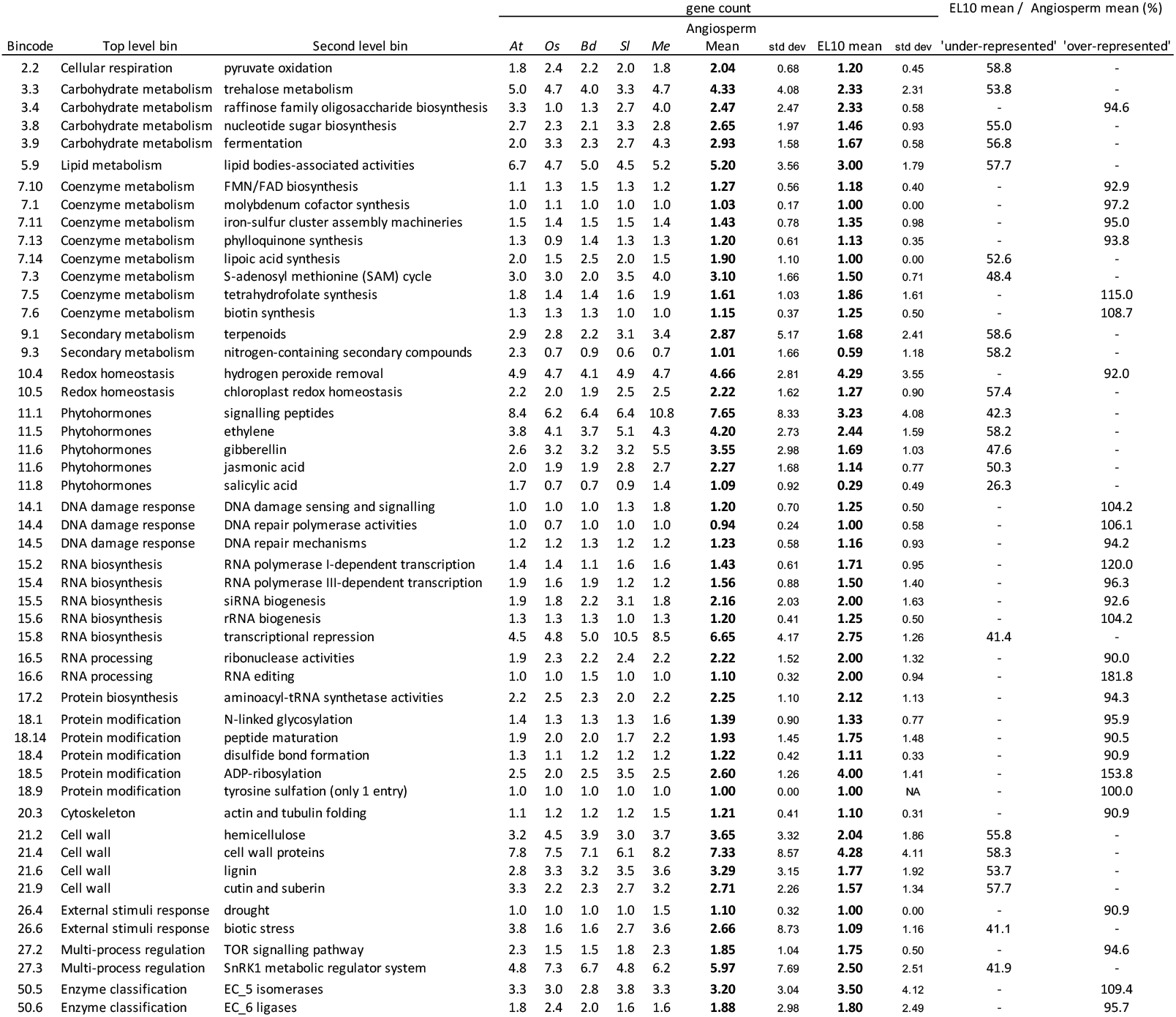
Comparison of over- and under-represented MapMan4 second-order functional classifications for EL10.1 and four other Tracheophytes [*Oryza sativa (Os) Brachypodium distachyon (Bd), Arabidopsis thaliana (At), Solanum lycopersicum (Sl), and Manihot esculenta (Me)*].

Phytohormone representation was low across all second-level categories, especially salicylic acid (Bincode 11.8). External stimuli response (Bincode 26) was rich in drought response but poor in biotic stress response genes. Multi-process regulation (e.g., integration of development with response-to-environment) was over-represented by the TOR signaling pathway (Bincode 27.2) and under-represented in the SnRK1 metabolic regulator system (Bincode 27.3). RNA biosynthesis (Bincode 15) was generally over-represented, however Bincode 15.8 (transcriptional repression) was greatly under-represented. Overall, 138 leaf bins were similar or over-represented and 447 were under-represented in EL10.1.

Transcription factor genes (Bincode 15.7) were under-represented overall. On average, there were ~10 fewer genes in EL10.1 than the average of five other angiosperms. Transcription factor genes with a >50 gene deficiency between the angiosperm average and EL10.1 included MADS box, NAC, MYB, and bHLH transcription factors (Table 12). Most of the transcription factor classes showing larger deficiency in copy number were members of larger gene families. Few transcription factor classes were equally- or over-represented, and most of these were from gene families characterized by lower copy number (Table 12). However, the FAR1 transcription factor class was abundant in EL10.1, and highly variable in the group of five other angiosperms (Table 12). It is likely that each of these differences in transcription factor copy number has potential to impact plant phenotype, development, and/ or response to the environment.

**Table 12:** Comparison of Transcription factor classes for EL10.1 and four other Tracheophytes [*Oryza sativa (Os) Brachypodium distachyon (Bd), Arabidopsis thaliana (At), Solanum lycopersicum (Sl), and Manihot esculenta (Me)*].

| Bincode | Description | gene count |  |  |  |  | Angiosperm |  | EL10 | Difference |
| --- | --- | --- | --- | --- | --- | --- | --- | --- | --- | --- |
|  |  | *At* | *Os* | *Bd* | *Sl* | *Me* | mean | stdev |  |  |
| 15.7.49 | FAR1 transcription factor | 18 | 89 | 142 | 44 | 36 | **77.8** | 48.8 | **106** | 28.25 |
| 15.7.3.7 | HB (Homeobox) superfamily.HOX-like transcription factor | 3 | 2 | 2 | 4 | 4 | **3.0** | 1.2 | **6** | 3.00 |
| 15.7.27 | ELF3 transcription factor | 2 | 2 | 1 | 4 | 2 | **2.3** | 1.3 | **3** | 0.75 |
| 15.7.4.3 | bZIP superfamily.bZIP19/23/24 transcription factor | 3 | 3 | 4 | 2 | 2 | **2.8** | 1.0 | **3** | 0.25 |
| 15.7.3.10 | HB (Homeobox) superfamily.SAWADEE transcription factor | 2 | 3 | 2 | 4 | 3 | **3.0** | 0.8 | **3** | 0.00 |
| 15.7.45 | HRT transcription factor | 3 | 1 | 1 | 1 | 1 | **1.0** | 0.0 | **1** | 0.00 |
| 15.7.5.4 | B3 superfamily.LAV-VAL transcription factor | 3 | 2 | 2 | 4 | 4 | **3.0** | 1.2 | **3** | 0.00 |
| 15.7.51 | SAP transcription factor | 1 | 0 | 0 | 2 | 2 | **1.0** | 1.2 | **1** | 0.00 |
| 15.7.53 | DPB3 transcription factor | 2 | 2 | 2 | 2 | 2 | **2.0** | 0.0 | **2** | 0.00 |
| 15.7.13 | HSF (heat shock) transcription factor | 24 | 25 | 24 | 27 | 32 | **27.0** | 3.6 | **16** | -11.00 |
| 15.7.46 | PLATZ transcription factor | 12 | 16 | 15 | 25 | 21 | **19.3** | 4.6 | **8** | -11.25 |
| 15.7.5.1 | B3 superfamily.ARF transcription factor | 23 | 24 | 24 | 25 | 29 | **25.5** | 2.4 | **14** | -11.50 |
| 15.7.3.13 | HB (Homeobox) superfamily.zf-HD transcription factor | 17 | 14 | 21 | 37 | 22 | **23.5** | 9.7 | **11** | -12.50 |
| 15.7.7.3 | AP2/ERF superfamily.AP2-type transcription factor | 18 | 22 | 24 | 24 | 30 | **25.0** | 3.5 | **12** | -13.00 |
| 15.7.16 | C3H zinc finger transcription factor | 57 | 52 | 49 | 76 | 68 | **61.3** | 12.9 | **48** | -13.25 |
| 15.7.19 | TCP transcription factor | 24 | 20 | 21 | 38 | 35 | **28.5** | 9.3 | **14** | -14.50 |
| 15.7.1.5 | C2C2 superfamily.DOF transcription factor | 36 | 30 | 29 | 36 | 44 | **34.8** | 6.9 | **20** | -14.75 |
| 15.7.1.3 | C2C2 superfamily.GATA transcription factor | 30 | 25 | 27 | 33 | 35 | **30.0** | 4.8 | **15** | -15.00 |
| 15.7.3.3 | HB (Homeobox) superfamily.HD-ZIP IV transcription factor | 18 | 12 | 17 | 49 | 13 | **22.8** | 17.6 | **6** | -16.75 |
| 15.7.39 | OFP transcription factor | 18 | 31 | 32 | 25 | 23 | **27.8** | 4.4 | **11** | -16.75 |
| 15.7.24 | AS2/LOB transcription factor | 43 | 36 | 28 | 52 | 56 | **43.0** | 13.2 | **26** | -17.00 |
| 15.7.7.2 | AP2/ERF superfamily.DREB-type transcription factor | 53 | 35 | 37 | 45 | 54 | **42.8** | 8.7 | **25** | -17.75 |
| 15.7.3.1 | HB (Homeobox) superfamily.HD-ZIP I/II transcription factor | 27 | 27 | 24 | 36 | 38 | **31.3** | 6.8 | **13** | -18.25 |
| 15.7.2.2 | MYB superfamily.MYB-related transcription factor | 81 | 72 | 64 | 81 | 85 | **75.5** | 9.4 | **55** | -20.50 |
| 15.7.2.3 | MYB superfamily.G2-like GARP transcription factor | 36 | 37 | 38 | 38 | 51 | **41.0** | 6.7 | **18** | -23.00 |
| 15.7.7.1 | AP2/ERF superfamily.ERF-type transcription factor | 63 | 60 | 52 | 87 | 97 | **74.0** | 21.4 | **38** | -36.00 |
| 15.7.12 | GRAS transcription factor | 34 | 57 | 63 | 53 | 78 | **62.8** | 11.0 | **26** | -36.75 |
| 15.7.4.1 | bZIP superfamily.bZIP transcription factor | 76 | 96 | 87 | 75 | 81 | **84.8** | 9.0 | **46** | -38.75 |
| 15.7.22 | WRKY transcription factor | 72 | 100 | 88 | 82 | 101 | **92.8** | 9.3 | **45** | -47.75 |
| 15.7.15 | C2H2 zinc finger transcription factor | 106 | 119 | 107 | 108 | 158 | **123.0** | 24.0 | **75** | -48.00 |
| 15.7.14 | MADS box transcription factor | 109 | 75 | 79 | 146 | 82 | **95.5** | 33.8 | **43** | -52.50 |
| 15.7.17 | NAC transcription factor | 113 | 136 | 135 | 102 | 111 | **121.0** | 17.1 | **61** | -60.00 |
| 15.7.2.1 | MYB superfamily.MYB transcription factor | 135 | 118 | 121 | 139 | 176 | **138.5** | 26.7 | **71** | -67.50 |
| 15.7.33 | bHLH transcription factor | 172 | 178 | 162 | 175 | 207 | **180.5** | 19.0 | **113** | -67.50 |
|  | mean of 91 classes | 21.5 | 21.4 | 21.4 | 23.8 | 25.6 | 23.1 | 5.2 | 13.7 | -9.4 |
|  | stdev of 91 classes | 32.0 | 33.1 | 33.0 | 33.9 | 38.1 | 33.7 | 7.8 | 21.3 | 15.6 |

### Genome size

Reported genome sizes (714 - 758 Mb; Arumuganathan and Earle 1991, derived from estimates for one plant each of table and sugar beet, respectively) and assembled genome sizes of sugar beet (~540.5 - 566.6 Mb, Table 2) may be explained by failure to assemble repetitive sequence arrays completely. To better assess genome size as a gauge of the completeness of assemblies in *Beta vulgaris*, an additional 50 independent cytometrically-determined nuclear DNA content estimates were obtained from four unrelated germplasm accessions; two traditional out-crossing progenies and two from progeny of deeply inbred accessions of EL10 and an inbred table beet derived from germplasm ‘W357B’. Nuclear DNA content estimates of these materials ranged from 633 Mb to 875 Mb, as estimated from at least four biological replicates from each accession (at least 20 from inbreds) with four technical replicates performed per biological replicate (Table 13). Overall, genome size between crop types was not statistically different (sugar beet, n = 120, mean = 729.0 Mb/1C, std. dev. = 51.2; table beet, n=80, mean = 742.3, std. dev. = 52.8; p = 0.079). Average genome size differences of each sugar beet accession were significantly different from one another (p < 0.001, means and dispersion values are presented in Table 13), and only the difference between sugar beet ‘5B sugar breeding population’ and Inbred Table beet was not significantly different than the other two sugar beet accessions. Inbreds showed a statistically-significant smaller average genome size (Table 13: inbreds, mean = 728.5 Mb/1C, out crossed, mean = 764.9 Mb/1C, p = 0.0002), and at least 2-fold higher variation than out-crossers (Table 13). The average cytometrically-determined genome size of all tested accessions was 734.3 Mb (stdev = 50.3 Mb). The smallest cytologically-estimated DNA content (633 Mb), coincidentally present in the progeny of EL10, closely approximated EL10’s optical map length of 628 Mb, and curiously, the average genome size of EL10’s progeny was 88 Mb larger than the assembled EL10.1 genome. Thus, average genome size appeared to increase over a single generation of selfing, and to an extent that reflected the size range observed within the species. This also implied that genome size also decreased at some point during the generation of these materials.

**Table 13:**
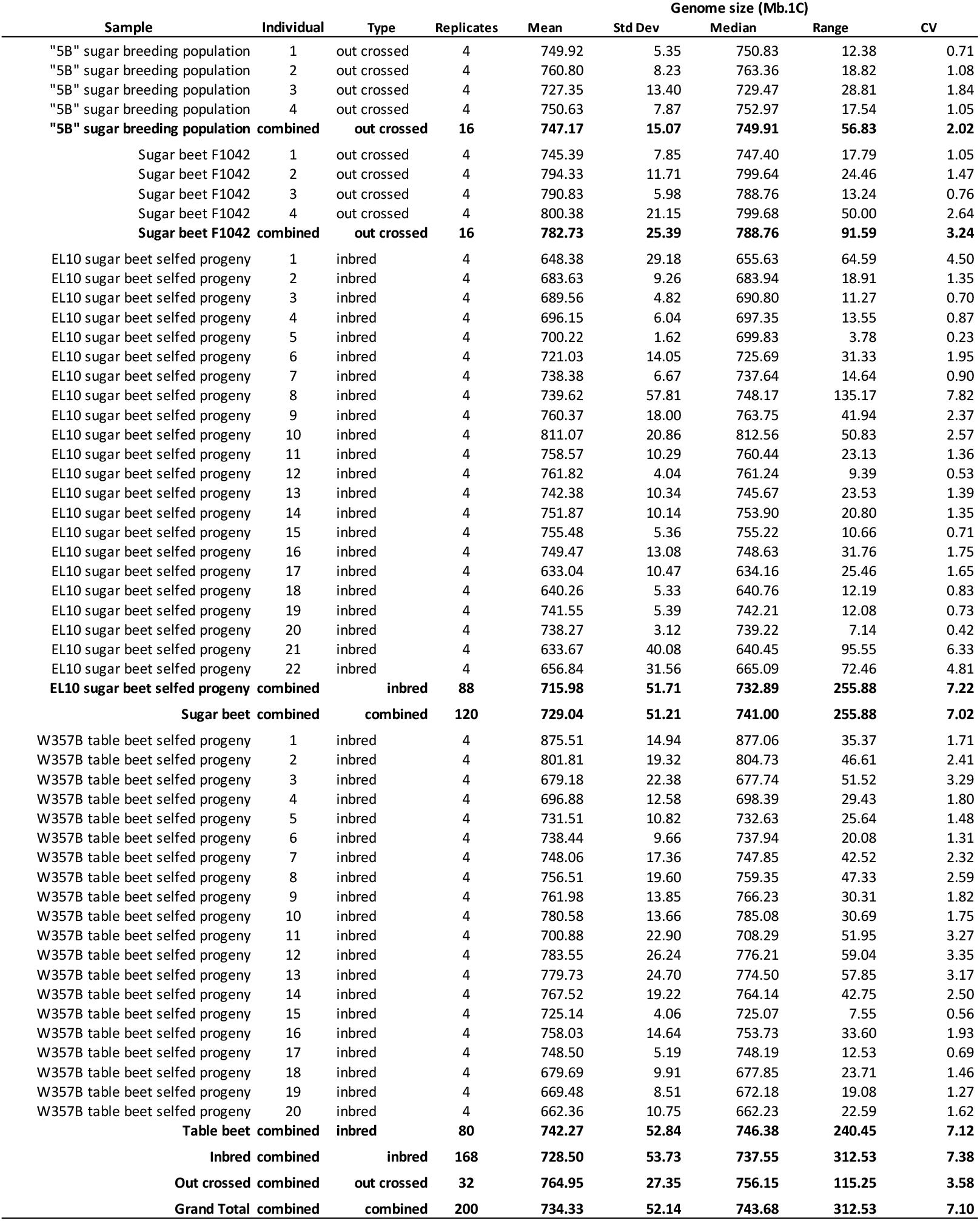
Beet genome size estimates obtained by flow cytometry.

### Repetitive element content estimation

Plant genomes are characterized by high repetitive sequence content, found either as tandem arrays or as multiple copies distributed throughout the genome (Bennetzen and Wang 2014). More than 180,000 named repetitive elements (as deduced by RepeatMasker) were placed on the EL10.1 assembly (Table 14). DNA class transposable elements were the most frequent (58.1%), which appears to be at odds with RefBeet (Dohm et al. 2014), and LTR elements the next most frequent class (36.0%) of annotated transposable elements (Table 7A). Numbers and types of LTR elements were estimated similarly using RepeatMasker and LTR_Retriever (Ou and Jiang 2018). However, distribution of the filtered high-confidence intact LTR_Retriever-predicted Gypsy and Copia elements (Table 14B) showed Copia elements generally more frequent towards the ends of Chromosomes and Gypsy elements biased towards centromeric regions (Figure 4).

**Figure 3:**
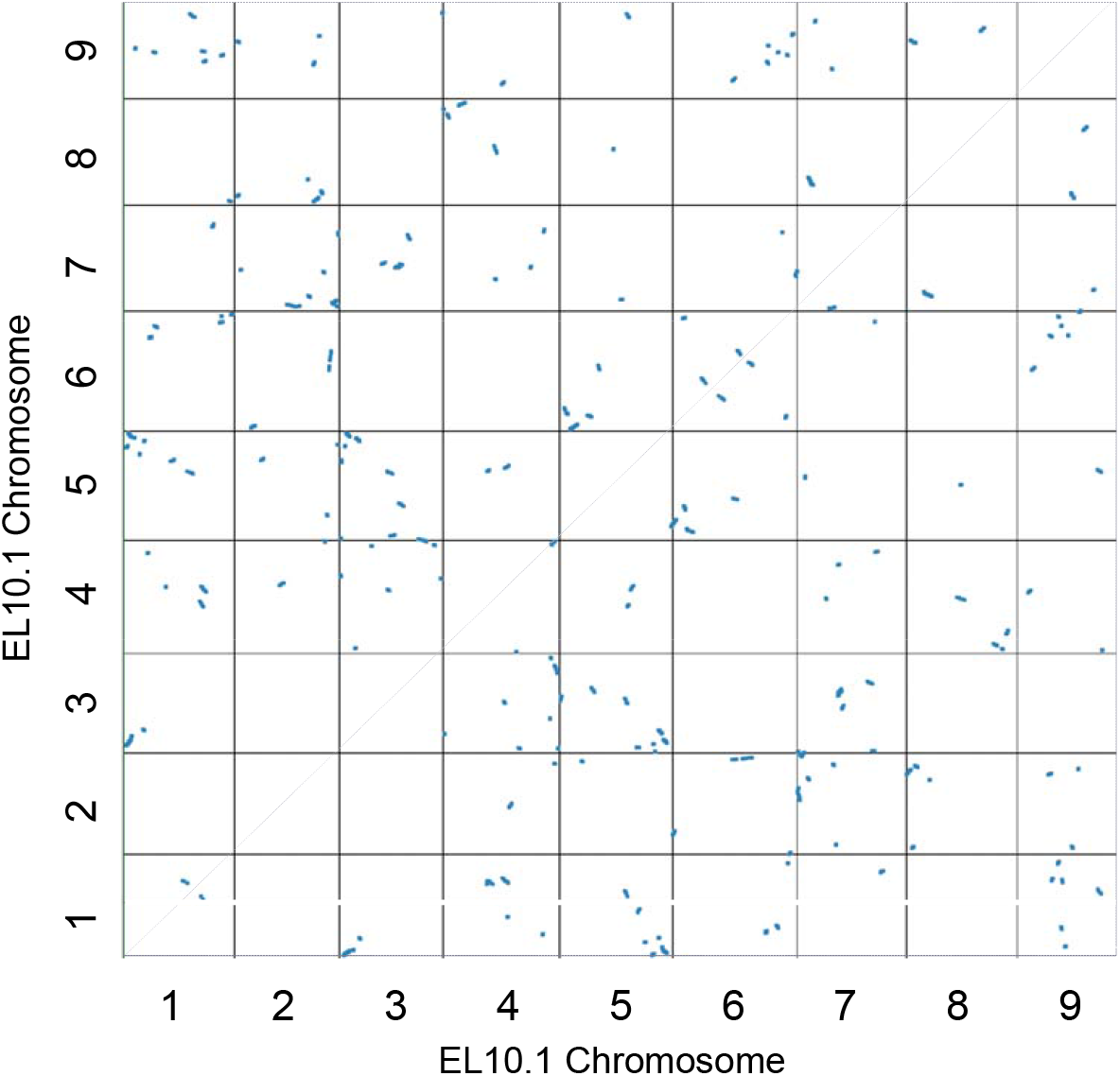
Self-synteny of EL10.1 Chromosomes against the EL10.1 predicted protein set.

**Figure 4:**
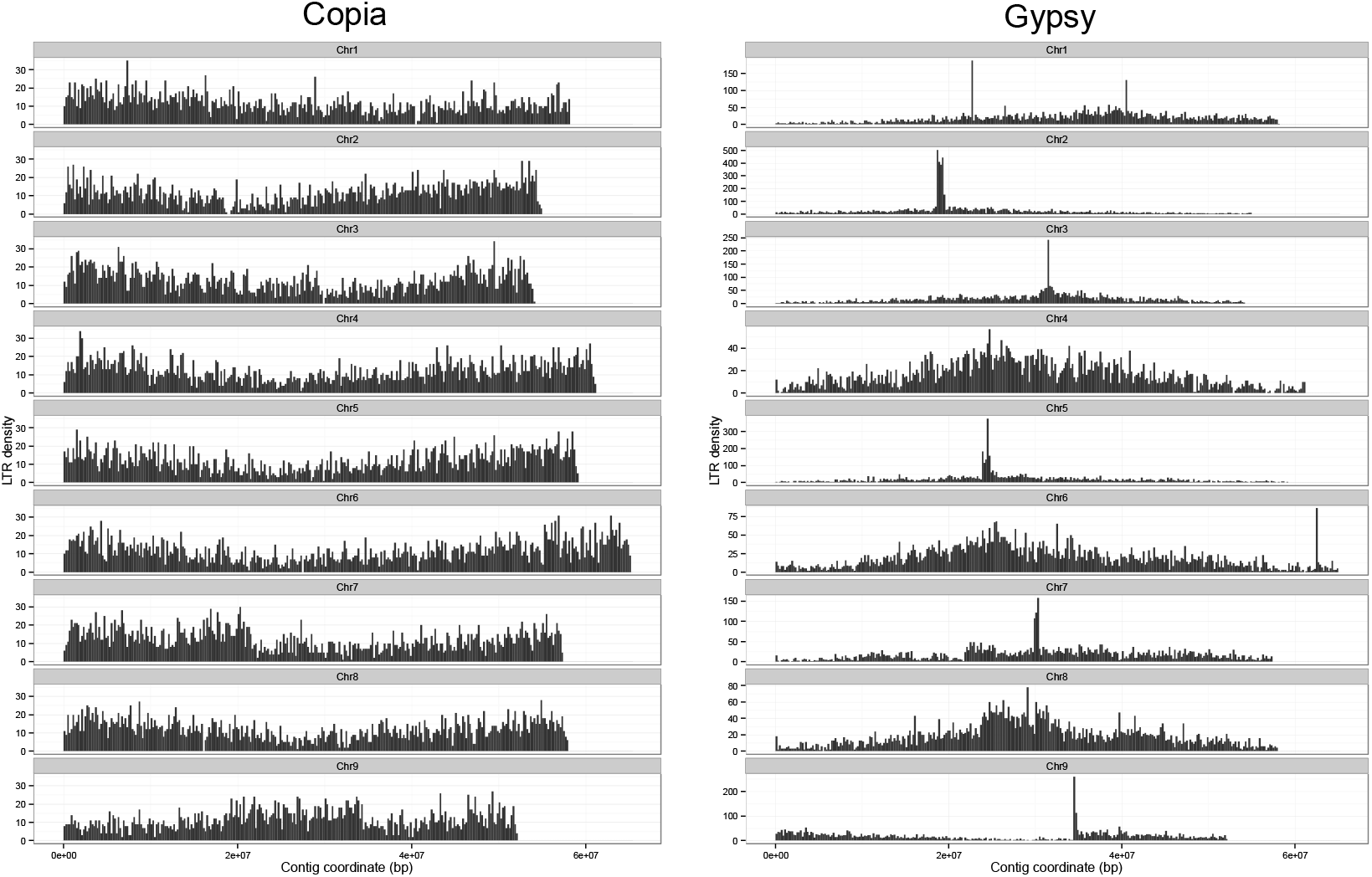
Distribution of LTR Copia and Gypsy retrotransposon elements across the EL10.1 Chromosomes.

**Table 14:** Frequency of Transposable Element (TE) classes in the EL10.1 genome assembly. A. RepeatMasker-derived annotations. B. Complete LTR retrotransposons. C. Interstitial repeat classes (from Kowar et al. 2016), D. Terminal repeat locations (from Dechyeva and Schmidt 2006) integrated with cytogenetic orientation (parentheses indicate reversed orientation relative to Paesold et al. 2012).

| Repeat Class | TE Type | Number in EL10.1 |
| --- | --- | --- |
| DNA | TcMar-Stowaway | 48,575 |
| DNA | En-Spm | 23,710 |
| DNA | hAT-Tip100 | 17,754 |
| DNA | MULE-MuDR | 6,473 |
| DNA | PIF-Harbinger | 3,878 |
| DNA | TcMar-Mogwai | 358 |
| DNA | MuDR | 87 |
| DNA | Maverick | 46 |
| LINE | L1 | 2,380 |
| LINE | RTE-BovB | 1,830 |
| LINE | L2 | 968 |
| LINE | DRE | 5 |
| LTR | Gypsy | 34,588 |
| LTR | Copia | 17,342 |
| LTR | undefined | 1,657 |
| LTR | Caulimovirus | 924 |
| LTR | ERV1 | 78 |
| LTR | Ngaro | 78 |
| LTR | Pao | 35 |
| MITEs | - | 4,481 |
| RC | Helitron | 3,869 |
| rRNA | - | 51 |
| Satellite | - | 5,587 |
| Simple | - | 2,227 |
| SINE | tRNA | 2,971 |
| SINE | 7SL | 679 |
| sum |  | 180,631 |
| mean |  | 6,947 |
| stdev |  | 12,050 |

**B: Complete LTRs (LTR\_Retriever)**
| Repeat Class | TE Type | Number in EL10.1 |
| --- | --- | --- |
| LTR | unknown | 999 |
| LTR | Gypsy | 1,030 |
| LTR | Copia | 574 |

**C: Major interstitial satellite sequences**
| Repeat Class | Name | Number in EL10.1 |
| --- | --- | --- |
| Gypsy CenH3 | Beetle7 | 1,937 |
| CenH3 | pBV_I | 4,029 |
| CenH3 | pBV_II | 3,863 |
| CenH3 | pBV_III | 4,387 |
| CenH3 | pBV_IV | 3,854 |
| CenH3 | pBV_V | 5,206 |
| CenH3 | pBV_VI | 5,090 |
| H3K9me2 | pEV1 | 10,463 |
| 5S rDNA | pXV1_5S | 107 |
| 35S rDNA | pZR1_18S | 199 |

**D: Major terminal satellite sequences (pAV34)**
| Chromosome orientation | Position in EL10.1 | Number in EL10.1 |
| --- | --- | --- |
| Chr1 - N | 34,591..49,546 | 38 |
| Chr1 | not found | - |
| Chr2-(S) | 1,010..7,486 | 21 |
| Chr2-(N) | 40,597,947..54,946,504 | 22 |
| Chr3 -N | 16,918..24,744 | 16 |
| Chr3-S | 54,068,612..54,100,447 | 83 |
| Chr4-(S) | 6..15,705 | 42 |
| Chr4-(N) | 39,426,832..61,140,574 | 95 |
| Chr5-(S) Scaffold 5 | 1,596,750..1,657,270 | 152 |
| Chr5 | 6,732,312..6,741,739 | 19 |
| Chr5 | 41,593,251..41,598,882 | 13 |
| Chr5-(N) | 59,198,215..59,224,585 | 71 |
| Chr6-(S) | 4..4,595 | 11 |
| Chr6-(N) | 6,5072,913..65,073,571 | 3 |
| Chr7-(S) | 1,002..15,434 | 45 |
| Chr7 | 8,866,608..8,879,321 | 24 |
| Chr7 | 21,826,741..21,836,557 | 26 |
| Chr8-(S) | 5..16,336 | 35 |
| Chr8-(N) | 43,367,644..57,938,902 | 101 |
| Chr9 | 25,507,827..26,781,523 | 3 |
| Chr9-(N) | 52,020,904..52,159,048 | 256 |

Repeats associated with centromeric histone variants have been characterized in beets (Kowar et al. 2016), and these consist of the Gypsy element Beetle7 as well the pBV class of major satellites (Table 14C). High-similarity Beetle7 sequences (90% identity over 1,000 nt or better) were located on all Chromosomes and eight of the Scaffolds. The 35S and 5S ribosomal RNA genes are also tandemly arrayed in beets (Paesold et al. 2012). The 35S arrays in EL10.1 were localized to Chromosome 2, as expected, and also to Scaffolds 7 and 19. The 5S array localized to Chromosome 4, also as expected, and to Scaffold 11. Only one canonical plant telomere array (TTTAGGG)_n_ greater than three tandem copies was found in the EL10.1 assembly, near the end of Scaffold 5. However, terminal repeat arrays defined by the major satellite class pAV (Dechyeva and Schmidt 2006) were found near the ends of most Chromosomes, except at one end each of Chromosomes 1, 5, 7, and 9 (Table 14D). pAV arrays were seen on each of these except Chromosome 1, where the South terminus appeared absent. Evidence suggests Chromosome 5 South is Scaffold 5, Chromosome 9 may have a pericentric inversion or an assembly artifact that misplaced Chromosome 9 South, and complex inversions in Chromosome 7 may have failed to accurately assemble the North terminal repeat region (these appeared to have been resolved in the EL10.2 assembly). Notably, interstitial pAV arrays were evident in both Chromosomes 5 and 7 (Table 14D).

Tandem repeats (unit length 500 nt or less assessed with Tandem Repeat Finder) were evenly spread across the EL10.1 assembly (Table 15), with an average of 630.4 repeats Mb^-1^ (stdev = 19.3) across Chromosomes, and similar for Scaffolds but with 25-fold higher variation (Mean 661.0 repeats Mb^-1^, stdev = 460.8). Shorter repeats were more frequent, and the most frequent size class was 21 nt (23,163 instances). Size classes of tandem repeats may reflect the predominant repeat unit size for centromeric sequence in a species (Melters et al. 2013), and for EL10.1, the most frequent repeat size above 100 nt was 160 nt (781 copies), followed by 170 nt (382 copies) (Figure 5). Relatively high numbers of repeats (67-134 copies) in the 314-325 nt repeat unit size range were evident, as might be consistent with a heterodimeric model of centromere repeats (Melters et al. 2013).

**Figure 5:**
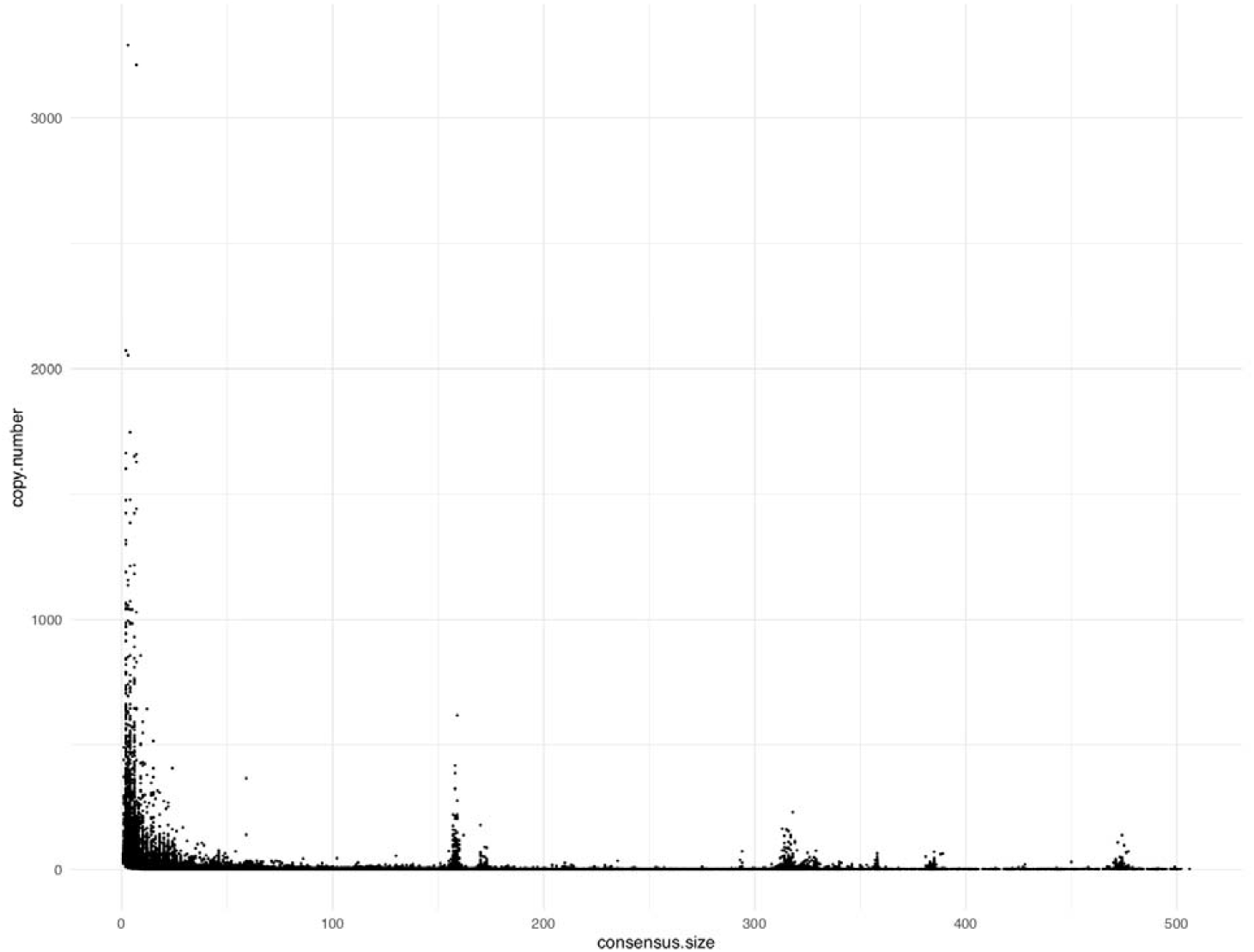
Copy number per consensus tandem repeat length in the EL10.1 genome assembly.

**Table 15:** Characteristics of tandem repeats in the EL10.1 genome assembly.

| Target | Length | Number of<br>Tandem<br>Repeats | Tandem<br>Repeats/ Mb | Mean Copy<br>Number | stdev | Median Copy<br>Number | Maximum<br>Copy<br>Number | Distinct<br>Types | Mean Percent<br>Match | Mean Indels |
| --- | --- | --- | --- | --- | --- | --- | --- | --- | --- | --- |
| Chromosome 1 | 58,086,001 | 36,834 | 634.1 | 7.8 | 35.7 | 2.8 | 3290.3 | 816 | 86.1 | 5.2 |
| Chromosome 2 | 54,971,872 | 37,041 | 673.8 | 7.5 | 27.3 | 2.9 | 2073.0 | 784 | 85.9 | 5.4 |
| Chromosome 3 | 54,100,447 | 33,183 | 613.4 | 7.8 | 24.1 | 2.8 | 1424.0 | 781 | 86.3 | 5.0 |
| Chromosome 4 | 61,163,185 | 38,116 | 623.2 | 7.9 | 26.4 | 2.9 | 1441.0 | 843 | 86.2 | 5.2 |
| Chromosome 5 | 59,224,585 | 37,876 | 639.5 | 7.8 | 23.3 | 2.8 | 1477.2 | 845 | 86.1 | 5.3 |
| Chromosome 6 | 65,096,967 | 39,448 | 606.0 | 8.2 | 28.3 | 2.9 | 1747.0 | 898 | 86.3 | 5.1 |
| Chromosome 7 | 57,353,724 | 36,137 | 630.1 | 7.5 | 23.3 | 2.8 | 1217.3 | 794 | 86.3 | 5.1 |
| Chromosome 8 | 57,938,902 | 36,560 | 631.0 | 7.8 | 22.2 | 2.9 | 1136.3 | 840 | 86.2 | 5.2 |
| Chromosome 9 | 52,180,088 | 32,462 | 622.1 | 8.1 | 26.3 | 2.8 | 1663.5 | 815 | 86.1 | 5.2 |
| Chromosome mean |  |  | 630.4 |  |  |  |  |  |  |  |
| Chromosome std dev |  |  | 19.3 |  |  |  |  |  |  |  |
| Scaffold 1 | 2,519,862 | 1,618 | 642.1 | 7.9 | 18.4 | 2.8 | 373.8 | 238 | 86.1 | 5.1 |
| Scaffold 2 | 2,420,327 | 1,529 | 631.7 | 8.9 | 21.0 | 2.8 | 383.0 | 261 | 86.4 | 5.1 |
| Scaffold 3 | 1,921,869 | 1,050 | 546.3 | 6.2 | 12.6 | 2.8 | 223.0 | 169 | 86.4 | 4.9 |
| Scaffold 4 | 1,802,165 | 1,173 | 650.9 | 8.9 | 30.8 | 2.9 | 661.5 | 201 | 85.9 | 5.4 |
| Scaffold 5 | 1,679,391 | 799 | 475.8 | 8.3 | 58.2 | 2.9 | 1628.4 | 159 | 86.5 | 4.8 |
| Scaffold 6 | 1,639,599 | 1,061 | 647.1 | 9.1 | 33.2 | 2.8 | 703.0 | 185 | 86.1 | 5.1 |
| Scaffold 7 | 1,327,247 | 705 | 531.2 | 7.2 | 16.4 | 3.0 | 269.0 | 142 | 86.9 | 4.9 |
| Scaffold 8 | 1,116,489 | 686 | 614.4 | 7.0 | 10.1 | 2.9 | 82.7 | 159 | 86.6 | 4.7 |
| Scaffold 9 | 912,454 | 420 | 460.3 | 9.0 | 59.2 | 2.6 | 1190.0 | 97 | 87.0 | 4.7 |
| Scaffold 10 | 511,739 | 290 | 566.7 | 10.2 | 23.3 | 2.9 | 230.2 | 98 | 86.7 | 5.2 |
| Scaffold 11 | 457,683 | 546 | 1193.0 | 4.1 | 9.8 | 2.9 | 196.0 | 66 | 84.9 | 8.0 |
| Scaffold 12 | 413,183 | 174 | 421.1 | 9.3 | 35.3 | 2.6 | 391.3 | 57 | 87.5 | 4.7 |
| Scaffold 13 | 378,582 | 244 | 644.5 | 9.6 | 31.9 | 2.8 | 384.8 | 77 | 86.5 | 5.0 |
| Scaffold 14 | 366,083 | 279 | 762.1 | 5.5 | 9.5 | 2.7 | 98.5 | 74 | 86.0 | 5.2 |
| Scaffold 15 | 344,704 | 109 | 316.2 | 8.2 | 14.9 | 2.7 | 88.8 | 49 | 89.0 | 4.1 |
| Scaffold 16 | 282,902 | 812 | 2870.3 | 3.3 | 3.6 | 3.0 | 70.9 | 47 | 84.5 | 9.8 |
| Scaffold 17 | 281,078 | 61 | 217.0 | 4.5 | 4.9 | 2.4 | 22.2 | 27 | 90.4 | 2.7 |
| Scaffold 18 | 278,623 | 108 | 387.6 | 7.2 | 12.2 | 2.6 | 89.0 | 48 | 88.2 | 4.0 |
| Scaffold 19 | 267,222 | 181 | 677.3 | 6.0 | 9.8 | 2.7 | 53.0 | 54 | 88.8 | 4.1 |
| Scaffold 20 | 264,226 | 131 | 495.8 | 6.9 | 11.5 | 3.1 | 93.0 | 55 | 85.5 | 6.0 |
| Scaffold 21 | 232,983 | 107 | 459.3 | 20.1 | 39.1 | 5.4 | 219.3 | 63 | 81.6 | 5.4 |
| Scaffold 22 | 195,802 | 96 | 490.3 | 6.3 | 10.3 | 2.7 | 61.5 | 40 | 88.7 | 4.6 |
| Scaffold 23 | 155,017 | 24 | 154.8 | 9.1 | 16.5 | 2.6 | 76.2 | 15 | 93.4 | 2.3 |
| Scaffold 24 | 136,997 | 260 | 1897.9 | 3.5 | 3.9 | 2.9 | 38.0 | 31 | 85.1 | 10.0 |
| Scaffold 25 | 135,273 | 39 | 288.3 | 6.0 | 8.5 | 3.0 | 46.0 | 29 | 84.4 | 6.5 |
| Scaffold 26 | 131,654 | 116 | 881.1 | 5.3 | 7.1 | 3.0 | 48.0 | 51 | 85.0 | 5.0 |
| Scaffold 27 | 130,158 | 55 | 422.6 | 11.0 | 15.9 | 3.1 | 57.0 | 38 | 86.1 | 6.9 |
| Scaffold 28 | 91,000 | 39 | 428.6 | 4.0 | 3.8 | 2.5 | 22.0 | 24 | 87.7 | 4.0 |
| Scaffold 29 | 14,771 | 6 | 406.2 | 6.7 | 10.0 | 2.9 | 27.0 | 5 | 92.7 | 2.0 |
| Scaffold 30 | 11,875 | 11 | 926.3 | 12.5 | 11.7 | 9.8 | 41.1 | 11 | 82.6 | 3.9 |
| overall mean |  |  | 661.0 |  |  |  |  |  |  |  |
| overall stdev |  |  | 460.8 |  |  |  |  |  |  |  |

Assembly continuity was accessed using the LTR Assembly Index (LAI) (Ou and Jiang 2018). After adjusting for the amplification time of LTR-RTs, the whole-genome LAI of the EL10 assembly was estimated to be 13.3, which is considered reference quality and improves upon the RefBeet assembly (LAI = 6.7) (Figure 6). Thus, the EL10.1 sugar beet genome assembly appeared to be largely complete with respect to repetitive element landmarks and assembled in a largely congruent fashion with respect to genetic markers.

**Figure 6:**
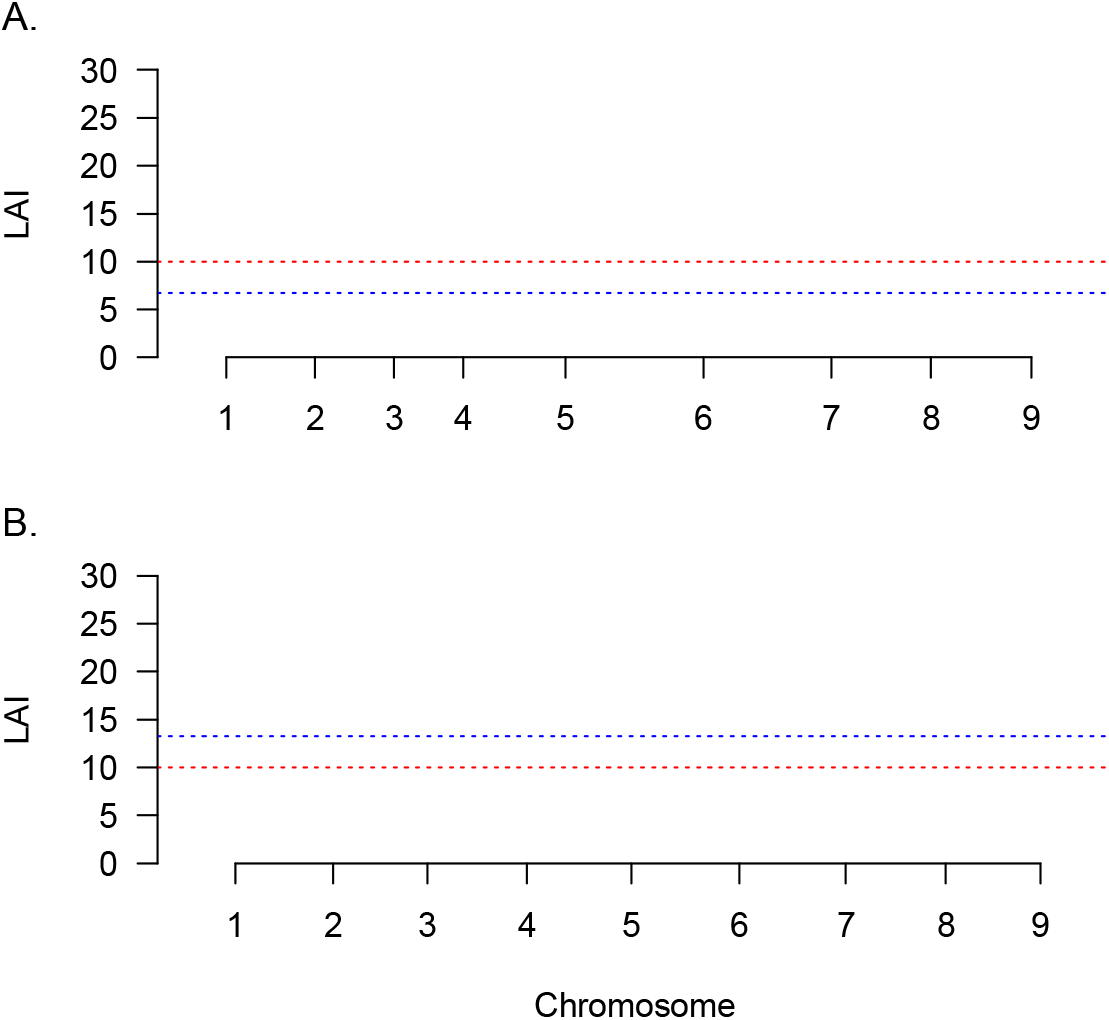
LTR Assembly Index (LAI) of the RefBeet assembly (A) and EL10 assembly (B) of the sugar beet genome. X-axes denote pseudochromosomes of the two assemblies. Each dot represents regional LAI in a 3 Mb window. Red-dotted lines indicate the LAI cutoff of the reference genome quality (LAI = 10). Blue-dotted lines indicate the mean LAI.

### Read count mapping

Read depth variation provided a means to compare accessions using readily available and deeper coverage short reads. Low variation in read depth suggests relatively even distribution of coverage across assembly coordinates, while higher variation suggests regions of low sequence complexity that may not have assembled in a consistent fashion, EL10 sugar beet genome - 13 perhaps contributing to differences in genome size between cytometry and assembly estimates. Five independent Illumina-derived read sets were read mapped to the EL10.1 genome assembly, one from EL10 and one each from four other sugar beet germplasms (including two EL10 relatives and two unrelated germplasms). Overall, more than 99.6% of EL10’s cleaned reads mapped to the EL10.1 assembly, with relatively even coverage (e.g. ~ 36 reads per assembled nucleotide), but Scaffold coverage was slightly less and the standard deviation was 22-fold higher. Similar results were evident in the other four germplasms (Table 16). There appeared no ‘degree-of-relatedness’ discrimination between disparate germplasm at this level of analysis, as EL10 relatives showed as much overall difference in read-depth variation as individuals drawn from unrelated populations (Figure 7).

**Figure 7:**
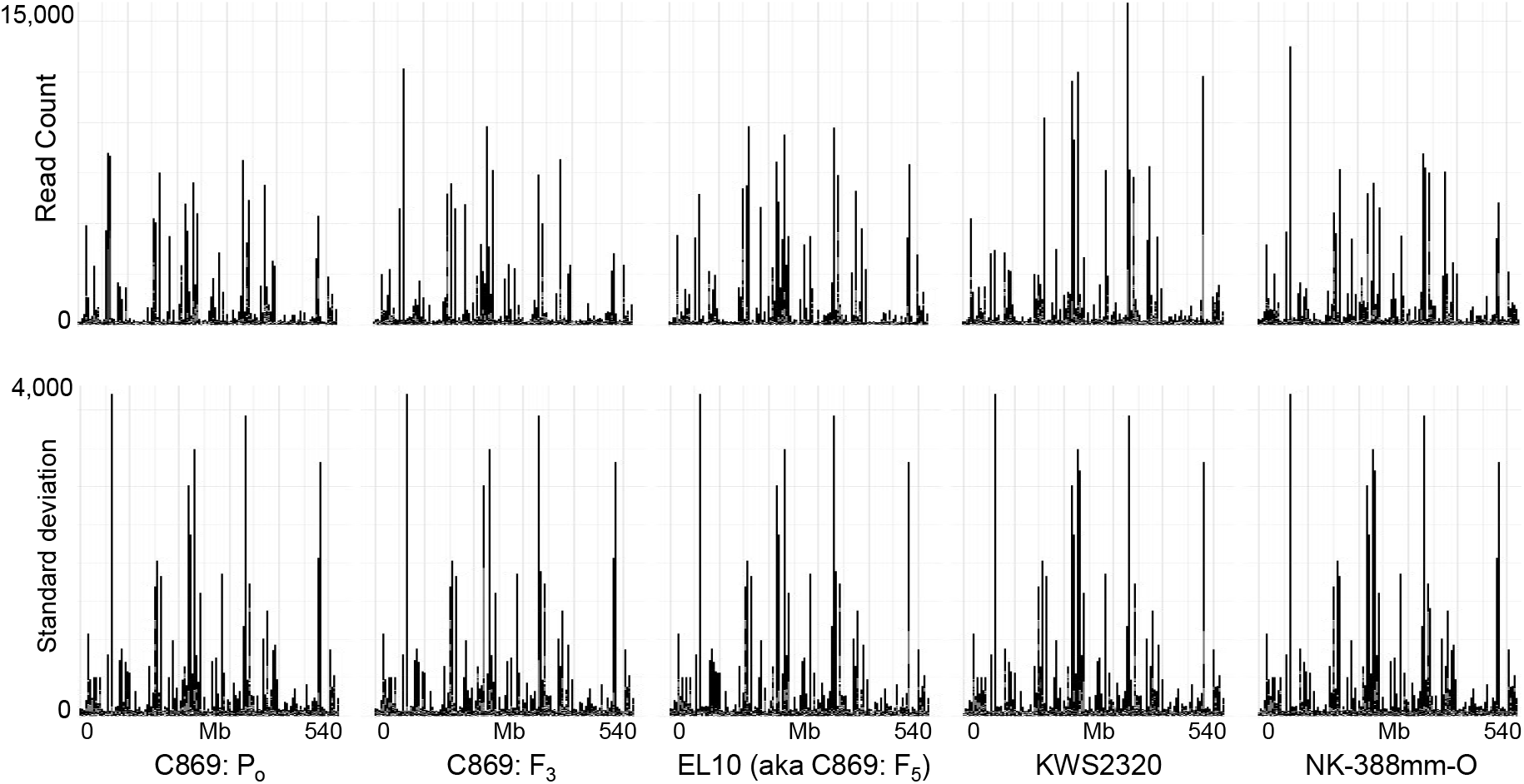
Read count mapping of short reads from EL10 and four other germplasms to the EL10.1 genome assembly and the standard deviation of reads mapped to each 5 kb window across the entire EL10.1 genome assembly.

**Table 16:**
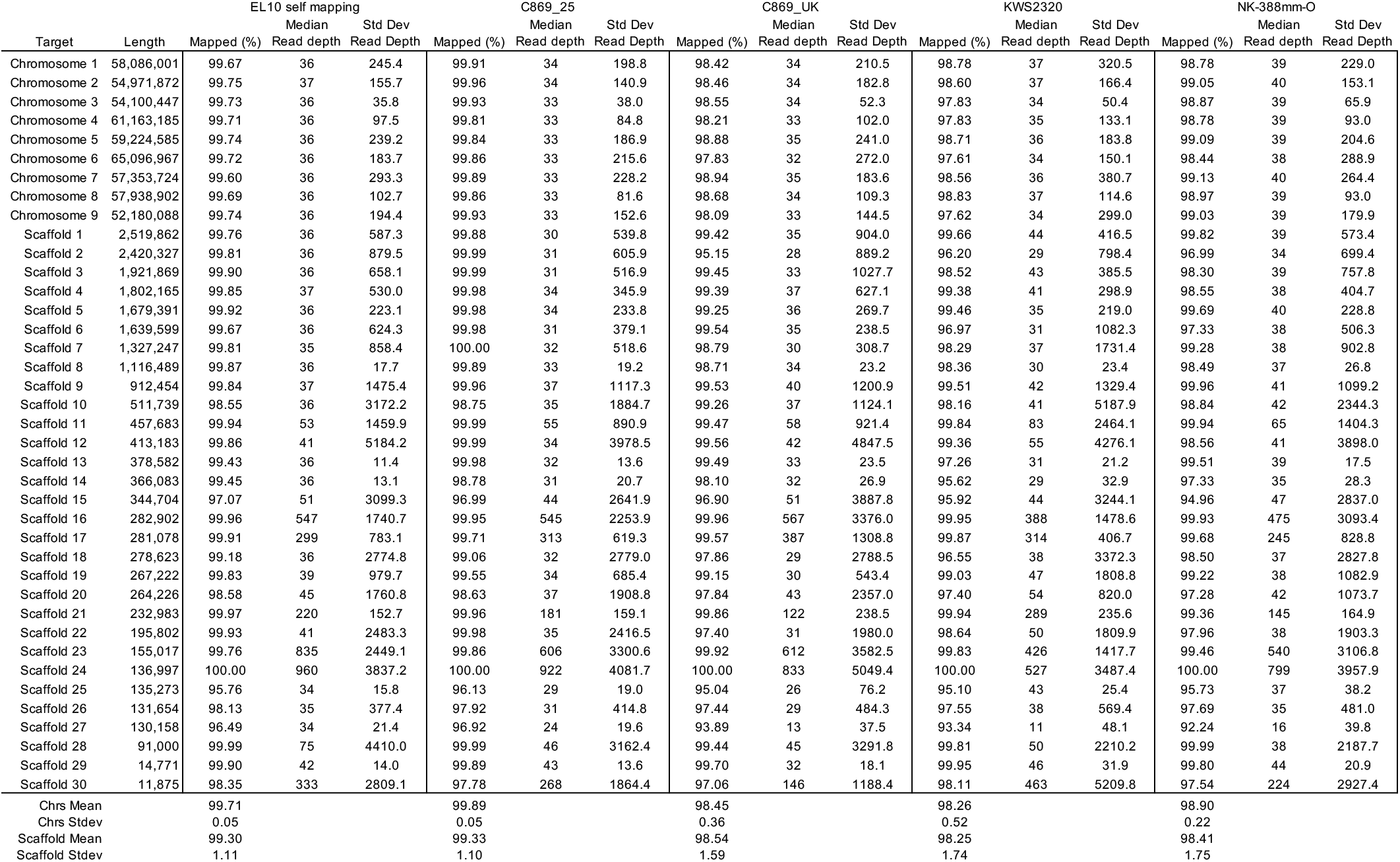
Read count mapping of short reads from EL10 and four other germplasms to the EL10.1 genome assembly.

High read-depth locations were localized using a conservative, computationally facile, and relatively crude sequence-independent approach. High read-depth locations were defined as a region of 5 kb with average per-base read mapping depth above 500 in one or more of the five tested germplasms (indexed from the lower nucleotide position of the EL10.1 assembly, Figure 8). This binning approach is conservative in the sense that most highly repetitive elements are shorter than the 5 kb window size used, but provided a computational advantage for an initial assessment whether changes in genome size could crudely be restricted to specific genomic bins, or were otherwise more or less independently distributed across the genome. The difference between C869_25 (i.e., the base genotype for EL10 and C869_UK) and each other accession flagged 47 such bins along Chromosome 1. Each flagged bin in each of the five germplasms occurred predominantly in the same places on Chromosome 1 (Figure 8). Most of these bins were occupied by Gypsy or Copia LTRs, however Bin 44,615,000 was occupied by chloroplast sequence (Sequence ID: KR230391.1) and Bins 8,100,000, 22,360,000, and 22,365,000 were occupied with mitochondrial sequences (Sequence ID: FP885845.1). It is not unusual to find plastid sequences within plant genomes (Pichersky et al. 1991), and plastid sequence read-depths are likely subject to external influences (e.g. plant growth and DNA isolation methods). The large differences in the remaining read-depth estimates at specific sites suggests that copy number changed since a last common ancestor. In the case of C869_UK and EL10, two generations of selfing had elapsed, and here differences were localized to 8 of the 47 bins on Chromosome 1 (Figure 8). These sites have the potential to contribute to intra-specific genome size variation. Further evaluation of such sites across the genome in a more precise sequencespecific fashion (e.g., not binned) may help deduce special features related to their lability and whether changes in genome size at this level of resolution have phenotypic effects.

**Figure 8:**
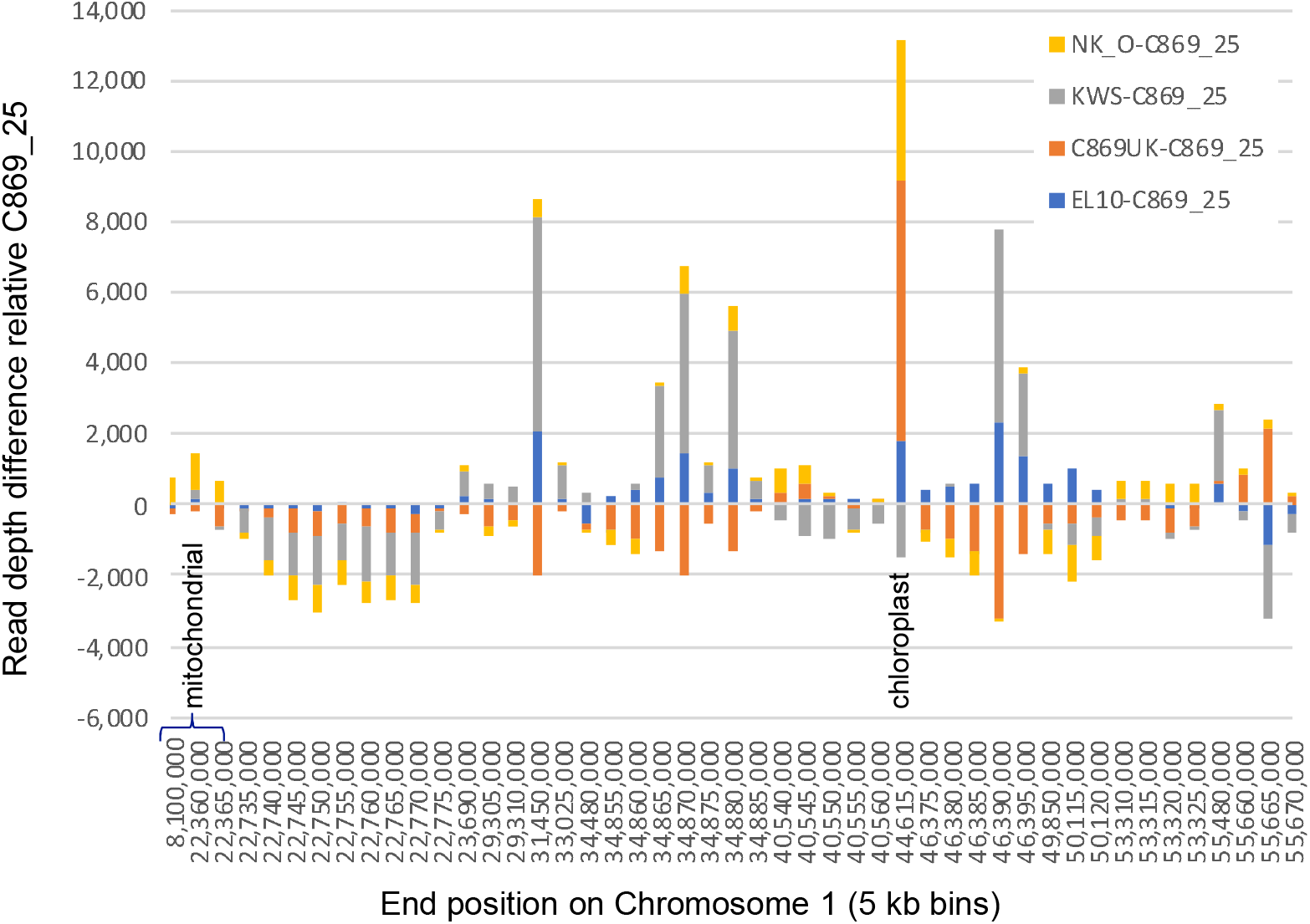
Distribution of high-copy number variant differences (>2000 copies per 5 kb window) between open pollinated population C869_25 and four inbred sugar beets across Chromosome 1 of the EL10.1 genome assembly.

### Broader synteny

Caryophyllales members spinach (*Spinacea oleracea*), grain amaranth (*Amaranthus hypochondriacus*), and quinoa (*Chenopodium quinoa*) have annotated genome assemblies that were used to compare with EL10.1 (Yang et al. 2016, Lightfoot et al. 2017, Jarvis et al. 2017, respectively; note that quinoa and amaranth are each amphidiploid). Chromosome 4 synteny appeared maintained in chromosome-sized blocks among Caryophyllales, as well as *Vitis vinefera* to a lesser extent, but not *Arabidopsis thaliana*, as outgroup representatives of the Rosids (Figure 9). Chromosome 1 synteny also appeared relatively conserved in chromosome-sized blocks among the Caryophyllales, with the exception of the spinach assembly version used here, which will likely improve in the future. Elements of Chromosomes 2, 6, and 9 were found in extended blocks in quinoa and amaranth, but also not spinach. Extended synteny for Chromosomes 5 and 8 were evident in quinoa but were not as extended in amaranth, while extended blocks for Chromosomes 3 and 7 were present in amaranth but not as well maintained in quinoa. Genome evolution within the Caryophyllales produced significant genomic variation in chromosome number, number of syntenic regions, and size of syntenic regions relative to beet (Table 17).

**Figure 9:**
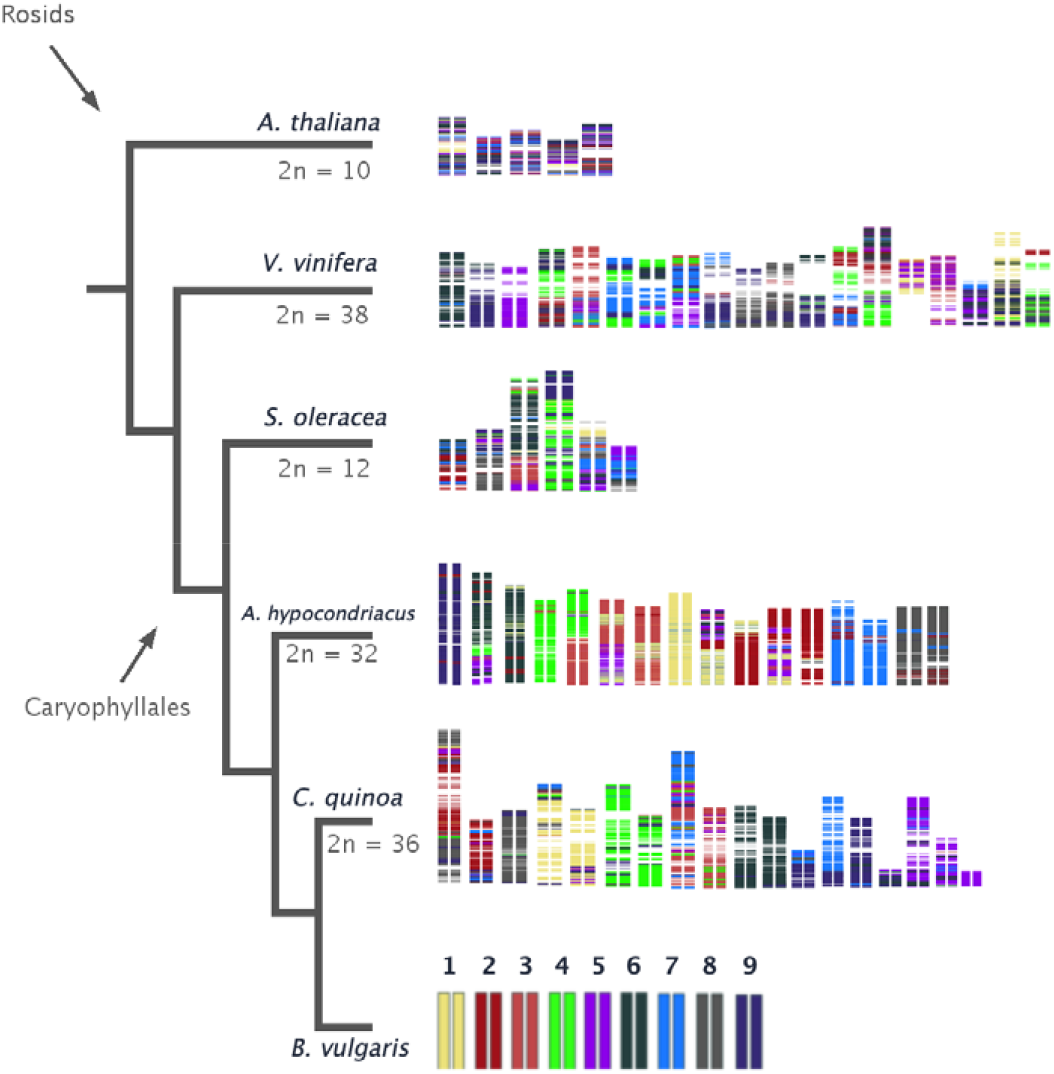
Visualization of syntenic blocks among Caryophyllales genomes relative to *B. vulgaris* EL10.1 Chromosomes compared with two representative Rosid species, color coded by EL10.1 Chromosome.

**Table 17:**
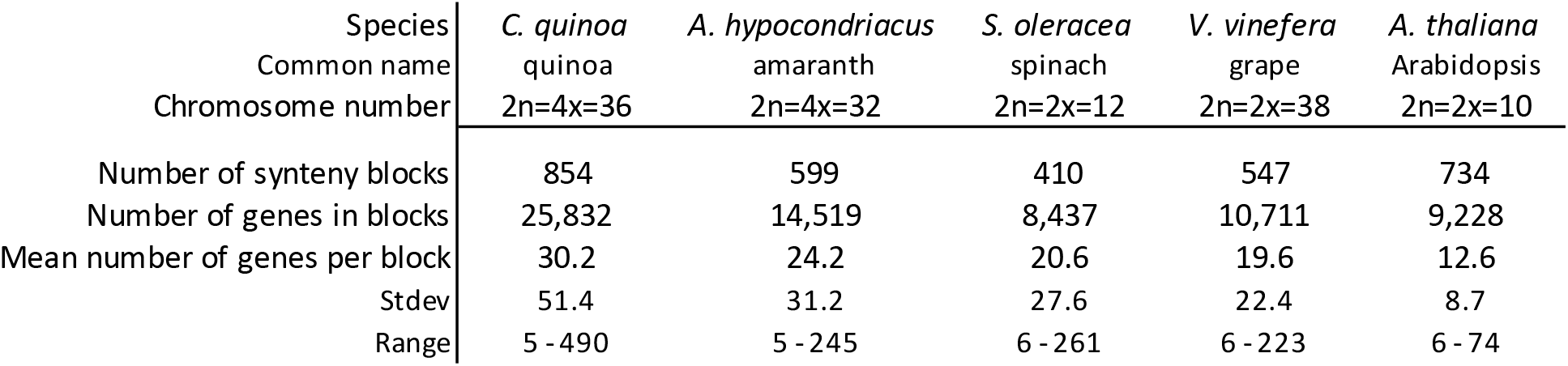
Proportion and metrics of synteny (co-linear blocks of MAKER beet gene predictions) shared among five species.

| Species | *C. quinoa* | *A. hypocondriacus* | *S. oleracea* | *V. vinefera* | *A. thaliana* |
| --- | --- | --- | --- | --- | --- |
| Common name | quinoa | amaranth | spinach | grape | Arabidopsis |
| Chromosome number | 2n=4x=36 | 2n=4x=32 | 2n=2x=12 | 2n=2x=38 | 2n=2x=10 |
| Number of synteny blocks | 854 | 599 | 410 | 547 | 734 |
| Number of genes in blocks | 25,832 | 14,519 | 8,437 | 10,711 | 9,228 |
| Mean number of genes per block | 30.2 | 24.2 | 20.6 | 19.6 | 12.6 |
| Stdev | 51.4 | 31.2 | 27.6 | 22.4 | 8.7 |
| Range | 5 - 490 | 5 - 245 | 6 - 261 | 6 - 223 | 6 - 74 |

## Discussion

A high-quality *de novo* assembly of the sugar beet genome was created. The EL10.1 assembly contains most of the ‘EL10’ genome organized into nine linkage groups plus 31 extra unplaced scaffolds. Most scaffolds contain predicted genes, and many were able to be placed in context of the larger chromosome-sized assemblies using genetic markers. Ends of chromosomes were captured to some degree, however additional work will be required to finish the EL10 genome assembly to exacting standards. Efforts to this end are underway and it should be noted that the EL10.2 assembly appears to resolve at least the major assembly-induced inversions evident of Chromosomes 7 and 9. Genome assembly is fraught with uncertainty. In most cases, there is no *a priori* information to gauge the completeness and correctness of an assembly. In this case, the fortuitous availability of a published sugar beet assembly (Dohm et al. 2014) allows for comparisons, however EL10 provides an independent perspective on the organization of the beet genome. The major difference between the two studies is that the sequencing technologies have improved to provide longer range scaffolding resulting in a substantially improved contiguity of sugar beet genome assembly. Such improvements also presumably better reflect copy number variations.

Long reads alone are currently insufficient for a high-quality assembly of a plant genome of moderate to large genome size. Beet might be considered a moderately-sized plant genome. The addition of an opto-physical map to long reads alone provided a ~10-fold reduction in the number of contigs, as well as set an upper bound for the size of the sequenced EL10 genome assembly (628 Mb). However, this also was insufficient to achieve chromosome-level assembly. Further addition of Hi-C data, where intact nuclei are cross-linked *in vivo* and where the native genome organization is presumably preserved, provided the means to map chromosome level associations. The sequential application of at least four independent technologies (i.e., short- and long-read sequencing, physical / optical maps, Hi-C chromatin conformation capture, and genetic maps) seems to have overcome many limitations spanning low-complexity regions of a genome over previous technologies in creating contiguous *de novo* genome assemblies of moderate to large plant genomes.

A reduction in gene copy number in beet (relative to annotated protein genes generated for comparative purposes, e.g., MapMan4) was observed. No clear evidence of gene copy number amplification was seen among the EL10.1 predicted protein set. Clear reductions were observed for transcription factors in particular, also observed by Dohm et al. (2014). Exceptions to the transcription factor reduction observed in EL10.1 included the FAR1 class of transcription factors, which may be anciently-derived from Mutator-like transposons and coopted in Arabidopsis for red-light perception and signaling (Hosoda et al. 2002, Mason et al. 2005). The role for this class of sequences remains unknown in beets, and copy number variation was high for FAR1 between the five other angiosperms considered. Discounting the occasional exception to lower overall gene copy number in beets, it may be suggestive of a basal gene copy number in dicots where beet numbers (or Caryophyllids in general) approximate a baseline condition, and other dicots have increased their copy numbers, as opposed to beets missing copies.

Genome size estimates of the cultivated beets examined here were quite variable, ranging from 633.0 to 875.5 MB per haploid genome. Genome size estimates of 21 wild *Beta vulgaris* spp. *maritima* genotypes from Portugal ranged from 660.1 to 753.1 MB (Castro et al. 2013), thus variability in genome size is known to occur in the species. The range of estimates was 2.6 times higher in the cultivated beets relative to the wild types. This was also observed relative to the breeding system of the cultigens, where the range in genome size among the out-crossers was 2.7 times lower than that of the inbreds (e.g., Table 13). Inbreeding *per se* may have effects on genome size in maize, including substantial loss of chromatin (Roessler et al. 2019). High variability in genome size between generations may be expected if copy-number variations were generated at each generation, as suggested by changes in genome size and read depth variation at specific loci. When during the life cycle such changes occur is not known, but presumably during a phase associated with DNA replication, and it is likely that transposable elements are at least partially involved (Whitney et al. 2010).

Variation in read-depth coverage may be useful for tracking genome size changes (Pucker 2019). Areas of high variation are intriguing from a chromosome biology and evolution perspective, as well as their potential effect on phenotype and on the origin of novel variation. It is no surprise that many plant genomes are large because of their highly repetitive nature, and many classes of repetitive elements are known to vary across kingdoms, often with little in common other than size, the fact they are repetitive, and characteristic footprints (target site duplications, terminal repeats, etc.) (Bennetzen and Wang 2014). Speciation seems to favor whole-scale sequence replacement of repeat elements while retaining their size, however interspecific amplified repeats seem to be present at low copy number in related genomes (Schmidt et al. 1991). Exactly how, and in particular when and what effects the efficiency, distribution, and specificity of divergent repeat amplification, is not as easy to investigate. Genome size reduction occurs rarely in plants, and improved read depth mapping approaches may be helpful in identifying additional examples and underlying mechanisms.

Two related, and two unrelated, germplasms contributed short-reads to the read depth differences observed, and are not necessarily representative breeding populations but rather have been crafted for genetic analyses. Beet is naturally a wind-pollinated out-crosser, which means that genetic diversity is partitioned within populations rather than between populations, as for inbreds. Each of the germplasms examined here, with the exception of C869_25, is highly inbred, using one of three different breeding methods. Both C869_UK and EL10 were derived from C869_25 through single seed descent, for three and five generations, respectively. RefBeet (aka KWS2320) was derived as a doubled haploid, and NK-388mm-O is a seed parent for hybrids inbred through conventional sib-mating (Taguchi 2014, Taguchi et al. 2019). The method used to generate the inbred seems not to relate to generation of read depth differences. Relatives of C869_25 showed as much difference in copy number as did the unrelated germplasm. However, each germplasm had a set of events specific to their own lineage, and others that were shared among two, three, or all germplasms. For instance, NK-388mm-O was enriched in depth at EL10.1 Chromosome 1 positions 53,310,000 Mb to 53,325,000 Mb, KWS2320 depauperate at positions 40,540,000 to 44,615,000, and C869_25 over represented from 22,735,000 to 22,7750,000. Responsible sequences underlying these regions have not yet been investigated, except where wide differences in chloroplast content, and less so mitochondria, were particularly rich in NK-388mm-O. We recognize the speculative nature of these interpretations, but they do generate testable hypotheses in a difficult to access arena of chromosome biology.

Exploration of synteny between species is accessible from a contiguous well-annotated genome sequence. For EL10.1, annotations were conservatively estimated from well curated plant gene resources, which likely improved confidence in assessing similarity between well known plant genes. Following the syntenic organization of such genes across phylogenetic groups showed that closely related species retained higher levels of synteny than more distantly related species, as expected. Also expected, was that recombination and schism of synteny blocks increased with increasing phylogenetic distance. Perhaps unexpected was differential synteny conservation by individual chromosomes. However, relatively few plant genomes are available that are highly contiguous, and this caveat limits interpreting results (for instance, the spinach genome is still under construction).

## Methods

### Plant material

USDA-ARS germplasm release EL10 (PI 689015) was derived by single seed descent from C869 (PI 628755) by self-pollinating over six generations. C869 is a biennial sugar beet conditioned by the self-fertile (*S^f^S^f^*) allele and is segregating for nuclear male sterility (*Aa*), with resistance to several diseases (Lewellen 2004). The initial selfing occurred from one self-fertile C869 CMS plant (EL-A013483) in 2002. Seed was field grown at the Bean and Beet Farm (Saginaw, MI) in 2005, roots were harvested, potted into fiber pots (5 L, Stock # ITMLFNP08090RBRD040TW, BFG Supply, Burton, OH), vernalized for 16 weeks, and grown in the greenhouse until flowering. Flowers were inspected for visible pollen, and when present, a #16 white grocery bag (Duro Bag, Novolex, Hartsville, SC) was placed over the bolting stem to effect self-pollination. Seed harvested from a single plant (EL-A018880) was considered the S1 generation, and subsequent generations were derived by single seed descent using field grown mother roots and selfing with the same methods. The S2 generation (EL-A022144) was obtained in 2007, and the S3 (EL-A025943) in 2010. Nine individuals of this population were genotyped with 69 SESVanderhave proprietary SNP markers evenly spaced across the beet linkage map, and a single homozygous individual (#17) of this population was sequenced for a preliminary assembly (named C869_UK here, McGrath et al. 2013). A sibling of this line (EL-A026195) with good field performance in the 2011 Michigan field (Saginaw Valley Research and Extension Center, SVREC, Richville, MI) was selfed in the same manner to yield the S4, while S5 (EL-A13-03870) and S6 generations were produced solely under greenhouse conditions in 2013 and 2015, respectively. Sixteen S6 individuals were genotyped with 24 SSR markers (McGrath et al. 2007), and six individuals (EL-A15-01096, EL-A15-01098, EL-A15-01099, EL-A15-01101, EL-A15-01102, and EL-A15-01103) were chosen as sequencing candidates based on marker homozygosity and similar growth habit and appearance, and pooled for long-read sequencing. One of these (EL-A15-01101) provided the sole tissue source for Illumina sequencing and nuclear DNA content estimation, and seed was named and released as EL10. Seed of EL10 was increased and deposited in the National Plant Germplasm System repository as a genetic stock (PI 689015).

Additional taxa were used, depending on the availability of materials. For the assessment of genome size, cytometric estimates were obtained from progeny of EL-A15-01101 whose genome was assembled here, advanced progeny of table beet W357B (a self-fertile parental line graciously provide by Dr. Irwin Goldman) which was inbred by single seed descent for five generations (accession EL-A1400766), an East Lansing open-pollinated self-sterile sugar beet breeding population (termed “5B”), and an open-pollinated USDA-ARS release used for a disease nursery check entry (F1042, PI 674103).

### Genome sequencing, assembly, and finishing

High molecular weight DNA for PacBio sequencing isolated nuclei using the HMW preparation protocols suitable for BAC library construction by Amplicon Express (Pullman, WA). PacBio RSII sequencing was performed at the Los Alamos National Laboratory (Los Alamos, NM), in 86 single-molecule, real-time cells using P6-C4 chemistry. PacBio reads greater than 6 kb were assembled with the Falcon Assembler (version 0.2.2), resulting in 938 primary contigs. Optical mapping was performed using the BioNano Irys sequential hybrid protocol with enzymes *Bss*SI and *Bsp*QI. For the EL10.1 assembly, scaffolding was accomplished using Proximity Guided Assembly (PGA) and Hi-C reads by Phase Genomics (Seattle, WA). Resulting scaffolds were polished and gap-filled using PBJelly, Arrow, and Pilon, following Bickhart et al. (2017). Briefly, PBJelly from PBSuite v15.8.24 was run using the Protocol.xml (https://gembox.cbcb.umd.edu/shared/Protocol.xml) with default parameters and minimum gap size set to 3 as: Jelly.py setup Protocol.xml -- minGap=3, Jelly.py mapping Protocol.xml, Jelly.py support Protocol.xml, Jelly.py extraction Protocol.xml, Jelly.py assembly Protocol.xml, and finally Jelly.py output Protocol.xml. Pilon v1.13 was run using --fix local bases and the is pipeline at: https://github.com/skoren/PilonGrid. Arrow v2.0.0 was run using the pipeline available at: https://github.com/skoren/ArrowGrid. And, Pilon v1.21 was run using --fix indels using the pipeline at: https://github.com/skoren/PilonGrid. For the EL10.2 assembly, 462 million Hi-C read pairs were input into the SBJ_80X_BN assembly (Table 2) and assembled via Chicago and Dovetail Hi-C technologies using the HiRise algorithm as described (Meyer and Kircher 2010, Putman et al. 2015). Bioinformatic manipulations during sequential assembly steps were performed by the respective organizations, and the ‘best’ assembly was then used as input for the next assembly step. Assembly metrics were assessed using assemblathon_stats.pl with default parameters (github.com/KorfLab/Assemblathon) (Earl et al. 2011).

### Whole-genome alignment

Whole-genome alignment of the EL10.1 assembly (as reference) and the RefBeet-1.2 assembly (as query) was conducted using modules from MUMmer v.4.0.0beta2. Initial alignments were created with the nucmer module, with options --mum -- minmatch 30 (uses only anchor matches that are unique in both the reference and the query, and sets the minimum length of a single exact match to 30 bp). The resulting delta alignment was filtered using the delta-filter module with options −1 -i 70 -l 5000 (to use only 1-to-1 alignments, with a minimum 70% sequence identity, and minimum alignment length of 5,000 bp). Summary reports were created using dnadiff, and plots were created from the filtered delta file using mummerplot with options --png --fat -r (with output image as png, and using layout sequences using fattest alignment only).

### Annotation

The EL10.1 assembly was annotated using the MAKER pipeline (Holt and Yandell 2011). The EL10.2 assembly has not been annotated. A custom repeat library for EL10 was created and used for repeat masking (Campbell et al. 2014). Protein and transcript evidence were used to aid gene model prediction. Protein evidence was obtained from the following species or databases: *Arabidopsis thaliana* proteins from Araport11 (Cheng et al. 2017), *Solanum lycopersicum* proteins from IPTG 2.4 (Fernandez-Pozo et al. 2015), *Populus trichocarpa* proteins from Phytozome genome v3.0 (Tuskan et al. 2006), and curated plant proteins from UniProt release 2017_03 (The UniProt Consortium 2017). Transcript evidence was derived from 25 RNA-seq read sets (BioProject PRJNA450098, Illumina 2500, 150 bp paired-end) using StringTie v1.3.3b (Pertea et al. 2015) and TransDecoder v5.0.1 (Haas & Papanicolaou et al., manuscript in prep. http://transdecoder.github.io).

Gene prediction programs AUGUSTUS (Stanke and Waack 2003) and SNAP (Korf 2004) were trained using the transcript sequences generated by StringTie (above), and both AUGUSTUS and SNAP were used to predicted gene models within the MAKER pipeline (Holt and Yandell 2011). When AUGUSTUS and SNAP predicted genes at the same locus, MAKER chose the gene model that was the most concordant with the transcript and protein evidence, and that model was retained at that locus. HMMER v 3.1 (Finn et al. 2011) was used to determine the presence of Pfam-A protein domains in the initial predicted protein sequences. Gene models supported either by protein or transcript evidence or by the presence of a Pfam domain were collected as high-quality gene models for the final genome annotation. Both transcript and protein sequences were searched against the SwissProt and UniRef databases using BLAST (Altschul et al. 1990). HMMER v3.1 (Finn et al. 2011) identified PfamA domains within predicted protein sequences. Signal peptide and transmembrane domains were predicted using SignalP v4.1 (Petersen et al. 2011) and TMHMM v2.0 (Krogh et al. 2001), respectively. Searches and predicted results were parsed and combined in the final functional annotation.

The online sequence functional classification and annotation tool Mercator4 ver. 2.0 (Schwacke et al. 2019) was supplied with the EL10.1 MAKER predicted protein fasta file using default settings. Four gene models were excluded from analysis due to their short length (<5 amino acids) (e.g. EL10Ac2g04429.1, EL10Ac8g20093.1, EL10Ac1g00658.1, EL10Ac7g16947.1). Comparisons were made with Mercator4-supplied representatives of the Tracheophyta (i.e. *Oryza sativa, Brachypodium distachyon, Arabidopsis thaliana, Solanum lycopersicum, and Manihot esculenta*).

### LTR annotation

*De novo* identification of intact LTR retrotransposons were performed using LTR_Retriever v1.6 with default parameters (Ou and Jiang 2018). The insertion time of each intact LTR-RT is estimated by LTR_retriever based on T = K/2μ where K is the divergence between an LTR pair and μ is the mutation rate of 1.3 x 10^-8^ per bp per year. Whole-genome LTR sequence annotations were achieved using the non-redundant LTR library generated by LTR_Retriever and RepeatMasker v4.0.0 (www.repeatmasker.org).

### LAI estimation

The assembly continuity of repeat space is accessed using the LTR Assembly Index (LAI) deployed in the LTR_retriever package (v1.6) (Ou and Jiang 2018). LAI was calculated based on either 3 Mb sliding windows or the whole assembly using raw_LAI = (Intact LTR-RT length * 100)/Total LTR-RT length. For the sliding window estimation, a step of 300 Kb was used (-step 300000 -window 3000000). The estimation of LAI was adjusted using the mean identity of LTR sequences in the genome based on all-versus-all BLAST.

### Tandem Repeats

Tandem Repeats Finder Program Version 4.09 was used to characterized tandemly dupicated sequences. using the default Alignment Parameters (e.g. match = 2, mismatch = 7, indels = 7, PM=80, PI=10, Minimum alignment score = 50, Maximum period size = 500) (Benson 1999).

### Self-synteny

CoGe SynMap (Lyons et al. 2008) was used, inputting *Beta vulgaris* (vEL10_1.0, id37197) and EL10.1 MAKER annotation gff files. Coding sequences were compared using LAST (Kielbasa et al. 2011) and DAGChainer (Haas et al. 2004) (with input settings Maximum distance between two matches = 20 genes, Minimum number of aligned genes = 5). Kn/Ks ratios (Yang 2007) were calculated using default parameters on CoGe (genomevolution.org/wiki/index.php/SynMap).

### Genome size variation

Four *Beta vulgaris* populations were evaluated for nuclear DNA content as described (Arumaguthan and Earle 1991). Briefly, young and healthy true leaf tissues from greenhouse grown seedlings were placed in between moist paper towels in zip-lock bags and shipped to the Flow Cytometry Lab at Benaroya Research Institute at Virginia Mason (Seattle, WA) for next day delivery. 50 mg of leaf tissue from each sample was finely chopped using a razor edge to release intact nuclei for flow cytometric analysis. Chicken erythrocyte nuclei (2.50 pg/2C) were used as an internal standard. A value of 978 Mb per pg was used for genome size conversion (Doležel et al. 2003). Statistical analyses were performed with JMP Pro version 14 (SAS, Cary, NC).

### Read count mapping

Reads from five Illumina paired-end sequencing datasets were trimmed and subsampled to produce sets of 25 GB for normalized mapping to the EL10.1 assembly. These were the single sequenced EL10 plant, a single plant two generations less inbred than EL10 (i.e., C869_UK), a pool of 25 individual from the parental population from which EL10 was derived (C869_25), the doubled haploid from which RefBeet was generated (KWS2320), and a single plant of a Japanese O-type breeding line (NK-388mm-O) (each accessible at NCBI BioProject PRJNA563463). Four samples of KWS2320 genomic reads (SRR869628, SRR869631, SRR869632, and SRR869633) were obtained from the NCBI SRA and pooled prior to filtering. FASTQ reads from the 5 mapping samples were filtered for a minimum FASTQ quality of 6 and minimum length of 80 bp after trimming. The reads that passed the filter were randomly subsampled to obtain 25 GB of reads per sample. Each pool of 25 GB was independently mapped to the EL10 assembly using BBMap v. 36.67 (Bushnell 2014). Read mapping was done with default parameters and kmer length = 13 with the addition of ‘local=t ‘to allow soft-clipping the ends of alignments and ‘ambiguous=random’ to randomly assign reads with multiple best matches among all best sites, to facilitate mapping of repetitive sequences evenly across the genome. For plotting read depth, 5 kb bins were created across each chromosome and the read coverage per base pair was calculated for each bin. The ‘basecov’ and ‘covstats’ outputs of BBMap were used to determine read depths and their standard deviations.

### Multispecies Synteny

The analysis of synteny was accomplished by plotting collinear blocks relative to beet chromosomes. Collinear blocks were defined using the program MCScanX using default recommendations (Wang et al. 2012). Protein sets for *A*. *thaliana, V. vinifera, S. oleraceae*, and *A. hypocondriacus* were downloaded from phytozome (https://phytozome.jgi.-doe.gov/pz/portal.html) with their corresponding gff files. Quinoa data were downloaded from chenopodiumdb (www.cbrc.kaust.edu.sa/chenopodiumdb/) and the *B. vulgaris* proteins and gff files were developed for this report.

### Accession Numbers

Sequence data from this article can be found in the EMBL/GenBank data libraries. The EL10 sugar beet whole genome project has been deposited in NCBI under the accession PCNB00000000. EL10.1 is version PCNB01000000. Associated NCBI database pointers are BioSample SAMN07736104, BioProject PRJNA413079; Assembly GCA_002917755.1, and WGS Project PCNB01. All raw reads used in EL10 genome assemblies are deposited in the short-read archive (SRA): Illumina reads SRR6305245; PacBio Reads SRR6301225; and Hi-C Library reads SRR10011257 (Phase Genomics) and SRR12507442 & SRR12507443 (Dovetail Genomics). BioNano Maps are located at SAMN08939661 (*Bsp*Q1) and SAMN08939667 (*Bss*S1). Read mapping accessions are deposited under BioProject PRJNA563463, and BioSamples SAMN12674955 (C869_UK), SAMN12674956 (C869_25), SAMN12674957 (NK-388mm-O). The EL10 genome assemblies and annotations can be viewed and downloaded via the CoGe Genome Browser available at genomevolution.org/coge/, both EL10.1 (Genome ID = 54615) and EL10.2 (Genome ID = 57232), and Phytozome only for EL10.1 (phytozome-next.jgi.doe.gov/info/Bvulgaris_EL10_1_0).

Genome browsing and file resources including transcript assemblies are available at sugarbeets.msu.edu. Transcript assemblies were constructed from root development and leaf RNA-seq reads derived from C869 (the EL10 progenitor) from 3 to 10 weeks post emergence (Trebbi and McGrath 2009) [3-week-old root (SRR10039097), 4-week-old root (SRR10039086), 5-week-old root (SRR10039081), 6-week-old root (SRR10039080), 7-week-old root (SRR10039079), 10-week-old root (SRR10039098), and mature leaf (SRR10037935)]. Also included were RNA-seq sets of 96 hr germinated seedlings from other germplasm germinated under aqueous stress conditions (McGrath et al. 2000), including 150 mM NaCl, 0.3% hydrogen peroxide, and biologically extreme temperatures (10 and 41 °C) (SRR10039075, SRR10039076, SRR10039077, SRR10039078, SRR10039082, SRR10039083, SRR10039084, SRR10039085, SRR10039087, SRR10039088, SRR10039089, SRR10039090, SRR10039091, SRR10039092, SRR10039093, SRR10039094, SRR10039095, and SRR10039096). The transcript assemblies are located at http://sugarbeets.msu.edu/data/EL10.1/.

## Supplemental Data files

To be selected from the contributed, at the discretion of the editors as appropriate.

## Acknowledgements

We thank Safa Alzohairy for isolating RNA from germinating sugar beet seedlings under various germination regimes. We also thank K. Arumuganathan, Director of the Flow Cytometry Lab of the Benaroya Research Institute at Virginia Mason, Seattle, WA for contracting the genome size estimates, and the staff at Dovetail Genomics (Santa Cruz, CA) for their assistance with the EL10.2 assembly. JMM: Funding provided by USDA-ARS CRIS 3635-21000-011-00D and the Beet Sugar Development Foundation, Denver, CO, USA. SK and AP were supported by the Intramural Research Program of the National Human Genome Research Institute, National Institutes of Health. Authors assert no conflicts of interest.

## Author Contributions

JMM, BT, EM-G, KD: Conceived and organized the work and wrote the manuscript; AF, PG, SO: Characterized EL10.1 assembly sequence organization; KD, HD, SJ: Created PacBio resources; JL, AH: Created BioNano resources; IL, SS, SK, AP: Conducted Hi-C assembly and finishing of EL10.1; AD, GW, SB, PS, KT: Applied proprietary genetic markers and materials to assess integrity of the EL10 assemblies; JW, TL, JP, KC: Provided MAKER gene annotations for EL10.1; AY, DF: Created RNA-seq transcriptome assemblies used in the EL10.1 annotation.

## Author emails

Mitch McGrath

Andrew Funk

Paul Galewski

Shujun Ou

Belinda Townsend

Karen Walston Davenport

Hajnalka Daligault

Shannon Lyn Johnson

Joyce Lee

Alex Hastie

Aude Darracq

Glenda Willems

Steve Barnes

Ivan Liachko

Shawn Sullivan

Sergey Koren

Adam Phillippy

Jie Wang

Tiffany Liu

Jane Pulman

Kevin Childs

Anastasia Yocum

Damian Fermin

Effie Mutasa-Gottgens

Piergiorgio Stevanato

Kazunori Taguchi

Kevin Dorn

